# Remodeling of a tripartite substrate-binding motif in the HD domain provides the mechanism for activation of CRISPR-Cas10 DNases

**DOI:** 10.1101/2022.11.07.515528

**Authors:** Zhenxiao Yu, Fang Wang, Qunxin She

## Abstract

Type III CRISPR systems are endowed with multiple immune activities, including target RNA cleavage, RNA-activated DNA cleavage and the cOA synthesis among which molecular mechanism remains elusive for the DNase. Here, DNase of LdCsm is investigated. Structural modeling revealed two HD loop segments. Cas10 mutants carrying either loop truncation or amino acid substitution in the HD domain are generated and characterized. We found each HD loop contains a substrate-binding site essential for its immunity. In fact, the substrate binding requires a tripartite motif composed of the two loop binding sites and the HD catalytic site. We demonstrate cognate target RNA (CTR) remodels the tripartite motif to activate the LdCsm DNase in two consecutive events: (a) it reduces the flexibility of LD-L1 and facilitates the simultaneous substrate binding at both loops, (b) the bound substrate is then propelled into the catalytic site via LD-L2 oscillation, driving the substrate into the catalytic site for DNA cleavage.

## Introduction

Clustered Regularly Interspaced Short Palindromic Repeat (CRISPR) and CRISPR-associated (Cas) systems code for the prokaryotic adaptive immunity that restricts invading genetic elements in a small RNA-guided fashion. The antiviral system is composed of two genetic units: CRISPR arrays consisting of invader-derived spacers interrupted by repetitive sequences and *cas* gene cassettes encoding Cas proteins. The immune response occurs in a microbe in three steps ^1-4^: (a) segments of invading nucleic acids are acquired and stored as spacers in CRISPR arrays, providing a record of previous encounters of invading genetic elements by the host, (b) biogenesis of crRNAs from the expression of CRISPR arrays, and (c) crRNAs and proteins synthesized from *cas* gene cassettes form ribonucleoprotein complexes (RNPs) in the host cell, which specifically recognize the re-occurring genetic elements and target them for destruction. To date, CRISPR-Cas systems are classified into two classes, six types which are further classified into more than forty subtypes ^5^. Among them, type III CRISPR systems belong to class 1 with Cas10 as the signature protein (also called CRISPR-Cas10). Six subtypes (III-A through III-F) are known, among which several III-A and III-B subtypes, which are also called Csm and Cmr systems, respectively, have been characterized.

Investigation of CRISPR-Cas10 systems demonstrates that they employ multiple immune activities to mediate antiviral defense. Initially, genetic study of the *Staphylococcus epidermidis* III-A (SeCsm) revealed that it mediates DNA interference in vivo ^6^, whereas purified *Pyrococcus furiosus* III-B (PfCmr) effector complexes specifically cleave RNA in vitro ^7^. Subsequently, it was demonstrated that the *Sulfolobus islandicus* III-B (SiCmr-α) mediates transcription-dependent DNA interference ^8^, exerting both DNA and RNA interference ^9^. In addition, *Streptococcus thermophilus* (St) Csm was also found to mediate RNA interference ^10^. By then, both subtypes were known to mediate dual DNA and RNA interference ^11, 12^.

Biochemical characterization of several CRISPR-Cas10 RNPs has identified active domains for each activity and their regulation. The large backbone subunits (Csm3/Cmr4) host the active site of the RNase of type III RNPs and they cleaves target RNAs in a 6-nt periodicity ^10, 13-15^. Cognate target RNA (CTR) carrying mismatches between its 3’-flanking sequence and the 5’-repeart handle of the corresponding crRNA activates unspecific ssDNA cleavage by the HD domain of Cas10 (Csm1/Cmr2) whereas noncognate target RNA (NTR) does not support that ^16-19^. The Cas10 Palm domains facilitate the synthesis of cyclic oligoadenylates (cOAs) and the activity is also subjected to the same regulation ^20-23^. cOAs function as a secondary signal that activates downstream effectors, most of which are nucleases ^24-30^, leading to cell dormancy or cell death ^25, 28^. More recently, we found that the *Lactobacillus delbrueckii* subsp. *bulgaricus* III-A (LdCsm) system, a unique CRISPR-Cas10 system, mainly relies on the DNase activity to mediate immune responses ^31^. Upon activation, the LdCsm effector cleaves transcription bubbles in vitro and the immune system indiscriminately cleaves accessible DNA substrates in the host cells, and mediating antiviral defense via abortive infection ^32^.

At the structural level, Cryo-EM structural analysis of RNPs of several type III systems has revealed the molecular mechanisms for target RNA cleavage in 6-nt periodicity and cOA synthesis ^33^. However, the mechanism involved in DNA cleavage remains elusive since only marginal differences are noted between their NTR-containing inactive forms and their CTR-containing active forms. Nevertheless, there are 1-2 flexible loops in the HD domain of III-A Csm structures ^34-36^. We compiled the reported structural data and found that there occurred apparent size changes for these loops in different conformations, suggesting CTR and NTR may differentially remodel these HD loops. We characterized the predicted loop segments in the LdCsm HD domain and revealed a tripartite substrate-binding motif with three binding sites distributed in the two HD loops and the HD active site. Therefore, CTR and NTR differentially remodel the tripartite motif such that CTR activates the DNase activity by facilitating the cooperation of the three substrate-binding sites, leading to DNA cleavage, whereas NTR facilitate their flexibility to prevent the occurrence of substrate binding at > 1 point, thereby inhibiting the CRISPR-Cas10 DNase.

## Results

### Structural modeling identified key factors that may regulate the LdCsm DNase

Structural modeling of the LdCsm RNP effector was conducted with high-resolution structures of StCsm, ToCsm and LlCsm as templates. These comparisons revealed that their HD domains are of two categories: StCsm and LlCsm possess two HD loop segments (HD-L1 and HD-L2) whereas ToCsm only has one (Fig. S1). LdCsm resembles more StCsm in structure (Table S1), thus carrying two predicted HD loop segments. The predicted HD-L1 segment stretches from N60 to S79, while the HD-L2 is from G92 to I114 (Fig. 1a∼b, Fig. S2c). In the simulated structure, HD-L1 occurs between the fourth and the fifth α-helix structures whereas HD-L2 is positioned after the fifth α-helix (Fig. S2a, 2c). Thus, the two loop segments are probably stabilized by different principles. HD-L1 is in the middle of two immediately adjacent α-helices, suggesting the boundaries of the loop could be redefined by remodeling of the two helices. For HD-L2, however, two conserved motifs may define its boundaries, including a GxDRR motif at one end and a pair of conserved amino acids (D109 and R121 for StCsm1) at the other end, the latter of which show polar interactions in the StCsm structure (Fig. S2a, S2c). Intriguingly, apo, NTR and CTR complexes of StCsm exhibit interesting differences in the HD region: (a) HD-L1 is highly flexible in the apo and NTR forms, but most of these flexible amino acid residues are in a more fixed position in the structure of the CTR-StCsm ternary complex. These data suggested that CTR and NTR differentially remodel the HD-L1 segment; (b) the flexibility of HD-L2 was kept more or less the same in all three conformations (Fig. S1, Fig. S2b), suggesting the conserved motifs at the ends of the loop segment firmly hold the HD-L2 segment and its flexibility could play an important role in DNA cleavage. Therefore, we characterized the two HD loops in the Cas10 (LdCsm1) subunit of LdCsm using genetic assays and biochemical characterization to test these hypotheses.

**Fig. 1.**
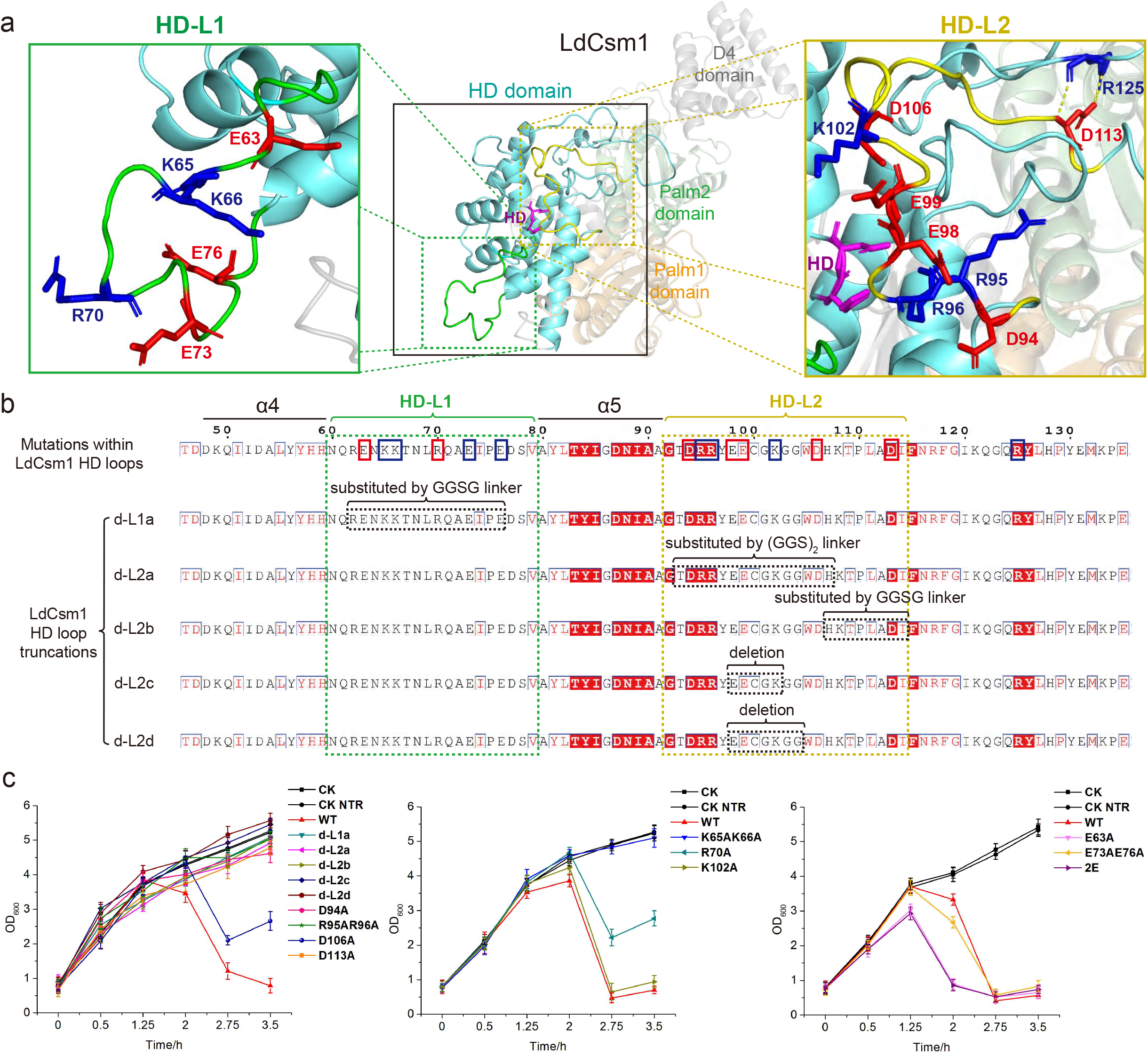
Identification of two HD loop segments in LdCsm and their functional characterization using the LdCsm interference assay. **a** Simulated structure of the LdCsm HD domain. The structure was obtained by using the SWISS-MODEL with the Cryo-EM structure of StCsm (PDB ID: 6ify) as the template. The HD active site is in purple; HD-L1 is colored green, while HD-L2 is colored yellow; positive charged amino acid residues in the loops are highlighted in blue; negative charged amino acids are in red. **b** Schematic of different mutations and truncations of LdCsm1 HD loops. HD-L1 and HD-L2 segments are in green and in yellow, respectively. Positively and negatively charged amino acid residues subjected to alanine substitution analysis are individually in blue and red solid boxes. Truncated loop segments that are either subjected to deletion or replaced with a short linker are represented with black dashed boxes. **c** Growth assay of the LdCsm DNase. Growth curves of test strains carrying HD-L1/L2 truncations or mutations. Wild type test strain (WT) contains pColE-Cas-S1 and p15A-CTR. Mutated LdCsm systems are indicated with the mutations in LdCsm1 they have carried, e.g., d-L1a represents the test strain contains pColE-Cas-S1-dL1a and p15A-CTR. Reference strain (CK) contains pColE-Cas-S1 and p15A-CK. NTR reference strain (CK NTR) carries pColE-Cas-S1 and p15A-NTR.

### Development of a new in vivo assay system to evaluate the LdCsm interference

In a previous work, we studied the in vivo DNA cleavage by LdCsm by analysis of plasmid mini-preps, and this revealed that LdCsm solely relies on the DNase to mediate immune responses. Nevertheless, the obtained results are unusual since nontarget plasmid is preferably targeted for degradation in the assay ^32^. Noticeably, the nontarget plasmid p15A-Cas-S1 has a low copy number whereas the target plasmid, pColE-CTR, is based on a high copy plasmid backbone (Fig. S3a). Thus, it is possible that their copy number difference could be attributed to the unusual outcome. To test that, backbones for construction of target plasmid and nontarget plasmid were exchanged to yield a new plasmid setup (Fig. S3b). Both plasmid setups were then tested experimentally for the assessment of LdCsm immunization using growth curves and plasmid loss rates. Test and reference strains for both assays were grown in LB broth to a mid-log phase. The resulting cultures were then divided into four equal portions, among which three were subjected to differential induction: (a) addition of L-arabinose to the medium (L-ara) to induce CTR production, (b) supplementing both IPTG and glucose (IPTG+Glu) to induce LdCsm synthesis but inhibits CTR production via catabolite repression, and (c) adding both IPTG and L-arabinose (IPTG+L-ara) to induce the synthesis of both LdCsm and CTR. Cultivation then continued for 4 h during which cell samples were taken for measuring OD_600_, determination of colony formation units (CFU) and plasmid loss rates.

For experiments with the reported plasmid setup, growth curves of the test strain resembled those of the reference in three of the four media, and only a slightly reduced cell mass was observed for the test strain in the L-ara medium in which the LdCsm synthesis is induced, relative to the reference (Fig. S3a). As a result, the onset of LdCsm immunization does not impair cell growth in the experiments of the reported assay. Determination of loss rates of both target and nontarget plasmids revealed that induction of the LdCsm immune responses only triggered the loss of p15A-Cas-S1, the nontarget plasmid (Fig. S3a), consistent with the reported data ^32^.

Conducting the experiments with the new setup of plasmids, however, yielded major differences in OD_600_ values and plasmid loss rates between the reference strain and the test strain. After attaining the peak value (∼3.5) at 1.25 h in the L-ara and IPTG+L-ara media, cell density (OD_600_) of the test strain rapidly decreased to (<1.0), and target plasmid (p15A-CTR) was readily lost from cells of the two CTR-induced cultures (Fig. S3b). This indicated that LdCsm immunization occurs along with the growth inhibition and loss of target plasmid in the induced cultures. In comparison, the reference strain grew similarly under the induced and non-induced conditions, and both target and nontarget plasmids were stably maintained. Determination of CFU of these cultures verified the above observations (Fig. S3b). These results indicated that the onset of the LdCsm immunity impairs the culture growth and facilitates the loss of target plasmid in the new plasmid setup. As a consequence, each of these physiological traits is useful for revealing the immune responses of LdCsm DNase in *E. coli*.

### Both loop segments in the Cas10 HD domain are essential for the LdCsm immune responses

Next, the new assay system was used to investigate whether the two HD loop segments could regulate the LdCsm DNase activity. Five loop truncations were designed, including (a) d-L1a (substitution of HD-L1 (R62∼E76) with GGSG), (b) d-L2a (substituting T93∼H107 of HD-L2 with (GGS)_2_), (c) d-L2b (substituting H107∼I114 of HD-L2 with GGSG), (d) L-L2c (deleting E98∼K102 in HD-L2), and (e) d-L2d (deleting E98∼G104 in HD-L2) (Fig. 1b). DNA fragments carrying the above truncations/substitutions were obtained by PCR and incorporated into the pColE vector to give the corresponding expression plasmids. These plasmids were introduced into *E. coli*, yielding test strains of Csm10^dL1a^, Csm10^dL2a^, Csm10^dL2b^ Csm10^dL2c^ and Csm10^dL2d^ (collectively called dL strains).

These dL strains were grown along with the wild-type LdCsm (WT) and the reference strains of the new assay. Cell samples were taken for determination of OD_600_, CFU and loss rates of p15A-CTR for each dL strain at specified time points. We found that induction of LdCsm synthesis in Csm10^dL1a^, Csm10^dL2a^, Csm10^dL2b^, Csm10^dL2c^ and Csm10^dL2d^ strains did not impair their growth, nor did the induction influence the maintenance of the target plasmid (Fig. 1c, Fig. S4a). These results are in full agreement with the data obtained with CK and in contrast to the data of WT, indicating that integrity and/or flexibility of each HD loop segment is essential for the activation of the LdCsm DNase.

### Charged amino acids in the loops differentially regulate the interference activity of LdCsm in *E. coli*

We noticed that both loop segments are rich in charged amino acids (Fig. S2a, 2c). Some of the charged amino acids are conserved, including D94, R95, R96 and D113 positioned at the two ends of HD-L2 whereas others are not. Non-conserved amino acid residues often have a relatively central position in the loops, including E63, K65, K66, R70, E73 and E76 in HD-L1 and E98, E99, and K102 in HD-L2 (Fig. S2c). These charged amino acids were mutated via alanine substitution and the resulting mutated LdCsm systems were analyzed using the developed genetic assay. Three distinct immune responses were found: (a) three mutations, including D94A, R95AR96A and D113A completely abolished the LdCsm immunity (Fig. 1c, Fig. S4a); (b) substitution of some positively charged amino acids impaired the antiviral immunity at different extents, including K65AK66A, R70A, and K102A (Fig. 1c, Fig. S4b), and (c) substitution of negatively charged amino acids in the loops facilitated the immune response and/or the level of plasmid clearance (Fig. 1c, Fig. S4c). Together, these results indicated that, while reducing positive charges in each loop impairs the LdCsm immunity, reducing the negative charges enhances it.

### The two HD loop segments are not equally important for the RNA-activated DNase of the LdCsm effector complex

To investigate how the DNase activity of LdCsm could be related to the immune responses, the WT and mutated LdCsm effector complexes carrying a truncation either in HD-L2 (d-L2c and d-L2d) or in HD-L1 (d-L1b: deletion of I74∼E76; d-L1c: deletion of A72∼E76 and d-L1d with both N68∼L69 and A72∼E76 deletions) were expressed in, and purified from *E. coli*. The resulting effectors of apparent homogeneity (Fig. S5a) were analyzed for DNA cleavage by PAGE. This revealed that loop 2 truncations (d-L2c and d-L2d) may have completely lost the DNase activity whereas three HD-L1 truncations are still active (Fig. S5b). Quenched-fluorescence DNA reporter assay further revealed d-L1b, d-L1c and d-L1d mutants retained, respectively, 23.5%, 8.5% and 6.9%, relative to the WT (Fig. 2a∼b). Clearly, HD-L2 is more important than Hd-L1 since any disruption to the loop integrity completely abolished the DNase activity. Furthermore, these results also suggested the DNase activity of the immune system can be modulated by changing the size of HD-L1.

**Fig. 2.**
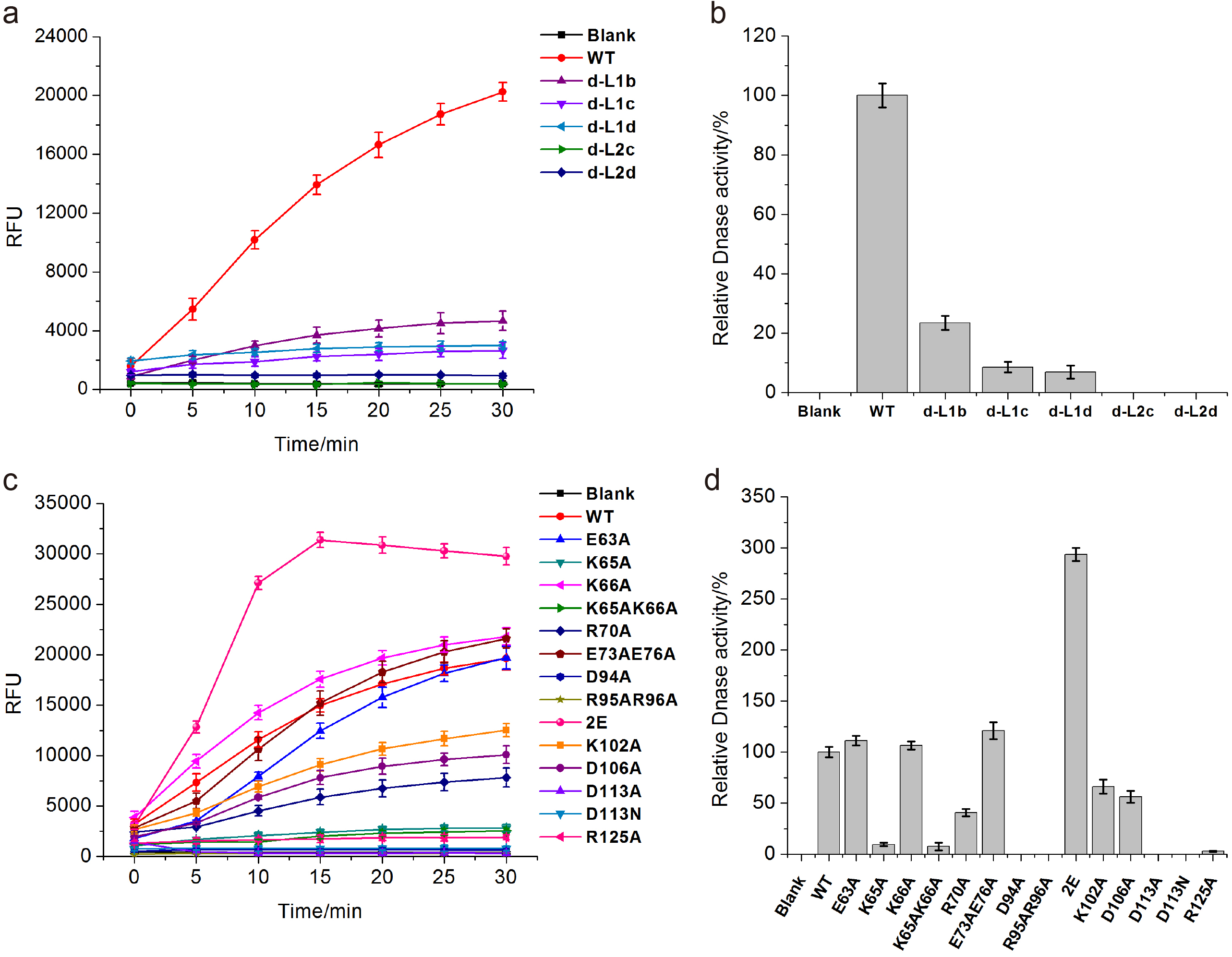
Fluorescence DNA reporter cleavage assay of the HD loop mutants of LdCsm. **a** Fluorescence DNA reporter cleavage assay of mutated LdCsm effectors carrying HD loop truncations in the LdCsm1 protein. **b** RFU increasing rates of the reactions in (a), the increasing rate of WT LdCsm is set as 100%. **c** Fluorescence DNA reporter cleavage assay of mutated LdCsm effectors carrying amino acid substitutions in the HD loops. **d** RFU increasing rates of the reactions in (c), the increasing rate of WT LdCsm is set as 100%. The cleavage assay was conducted with 50 nM effector complex, 1μM Fluorescent quenched ssDNA reporters and 500 nM CTR in the reaction buffer in a microplate reader at 37°C.

LdCsm effector complexes carrying E63A, K65AK66A, R70A, E73AE76A, D94A, R95AR96A, 2E, K102A, D106A or D113A substitution were also purified to an apparent homogeneity (Fig. S5a) and characterized as described above. We found that several LdCsm effectors are active in the presence of CTR but none is active in the presence of NTR (Fig. S5c), suggesting that the mechanisms of CTR activation and NTR inhibition are strictly maintained in all mutants. Quantification revealed that the DNase activity of LdCsm effectors carrying E73AE76A (in HD-L1) or 2E (in HD-L2) substitutions is about 1.2- and 2.9-fold higher than the WT type whereas K65AK66A, K65A, R70A, K102A and D106A substitution mutants exhibit residual activities of 7%, 9%, 36%, 60% and 56% individually (Fig. 2c∼d). Together, this suggested that K65 may serve as the main contact site for substrate binding in HD-L1. Interestingly, the two predicted loop anchors in HD-L2, including the conserved GxDRR at one end, and D113 and R125 at the other end, are particularly important since their mutations either yielded a DNase-dead LdCsm effector, including D94A, R95AR96A and D113A, or a strongly impaired DNase as for R125 (Fig. 2c∼d). These results suggested that HD-L2 could have another function in addition to substrate binding, in which its loop anchors play an essential role.

### Non-conserved negatively charged amino acids in the loops contribute to the interference by modulating substrate binding

We noticed that seed cultures of mutated LdCsm strains carrying E63A or E98AE99A (2E) substitution did not grow so well in LB medium; their OD_600_ values only reached 1.5∼1.8, compared to 3.1∼3.3 for the wild-type and other seed cultures (Table S2). This suggested that these mutations could have changed the threshold of the LdCsm system to respond invading RNA signals. As a result, the residual expression of CTR from the *araBAD* promoter in LB can attain the threshold to trigger the immunization of the mutated system. To study that, strains of the wild-type, or E63A or 2E mutated LdCsm (Fig. 3) carrying each of p15A-CK, p15A-CTR and p15A-NTR plasmids were grown in LB plus glucose to generate seed cultures. Interestingly, these seed cultures grew equally well in the glucose-containing medium, suggesting the presence of glucose reduces the CTR expression to a level below the new threshold. Then, interference assay was conducted for each LdCsm system in 3 different media (p15A-CTR in LB, LB+0.5%Glu and LB+1%Glu) with two references (p15A-CK and p15A-NTR). We found, whereas the WT LdCsm grew equally well in both glucose-free and glucose-containing media, greatly reduced cell mass (measured as OD_600_) was observed for both mutated LdCsm systems in the absence of glucose (Fig. 3). These data demonstrate that the two mutated LdCsm systems have gained the capability of responding to a lower level of CTR to activate their antiviral immunity, compared to the wild-type form.

**Fig. 3.**
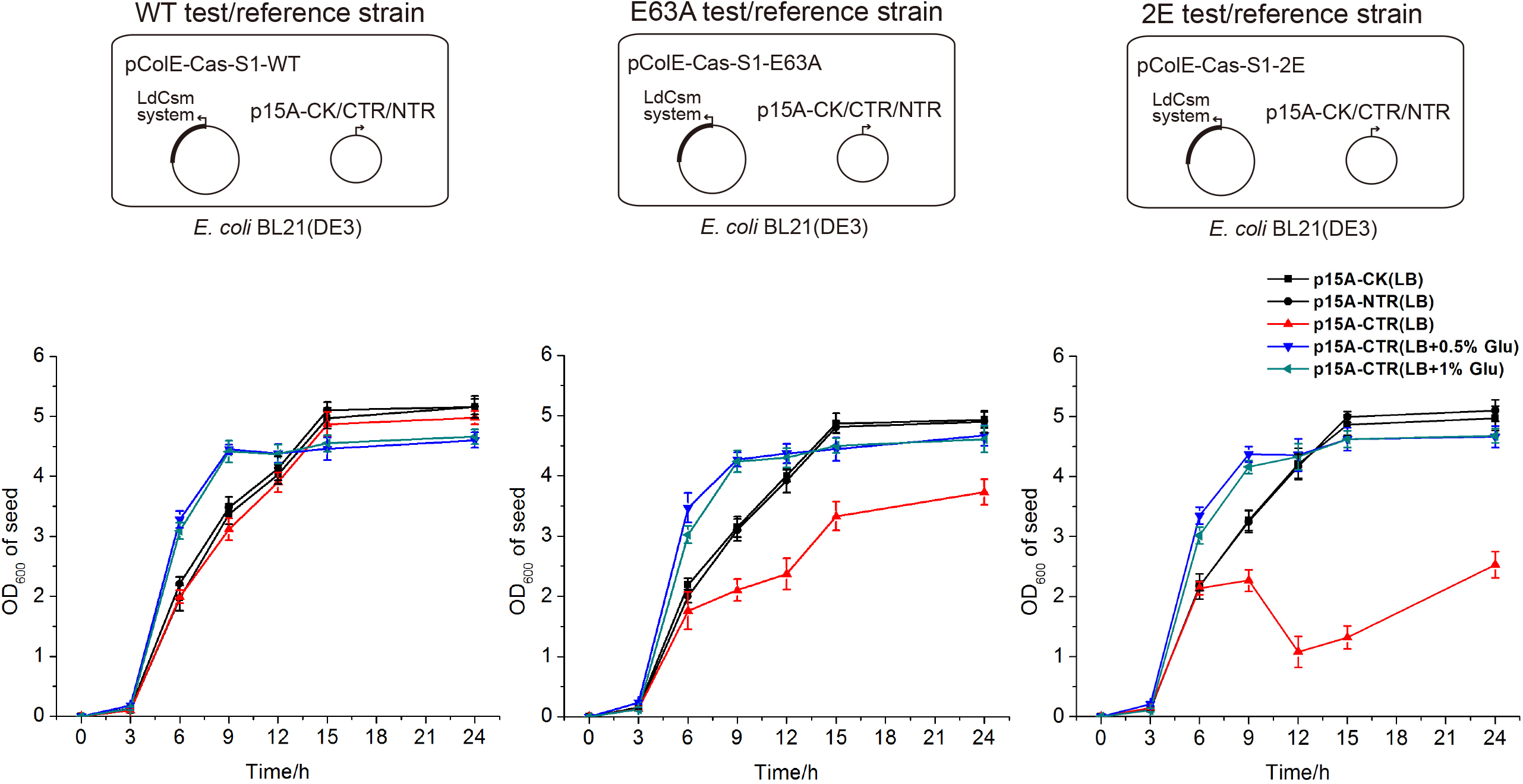
Reduction of negative charge in the HD loop segments enhances the LdCsm interference. Upper panels: plasmids employed in the three sets of experiments, which are the combinations of an LdCsm expression plasmid (WT, E63A or 2E (E98AE99A)) and a target RNA-expressing plasmid (CTR, NTR or CK). Lower panels: Growth curves of the three sets of experiment. Single colonies inoculated to 5ml LB containing Amp (100μg/ml) plus Kan (50μg/ml) or LB containing Amp plus Kan plus Glucose (0.5%/1%) and cultivated at 37°C, 220rpm. OD_600_ was measured for each culture during inoculation and plotted against incubation time.

To investigate if the elevated sensitivity of the mutated LdCsm effectors could be related to an enhanced substrate affinity, kinetics study was employed to investigate the ssDNA substrate affinity of three active LdCsm effectors, including LdCsm-dCsm3, LdCsm-dCsm3-E63A and LdCsm-dCsm3-2E. Noticeably, their dCsm3 variants were employed in the kinetic assay since inactivation of the Csm3 RNase strongly elevates the DNase of the LdCsm system ^31^. ssDNA cleavage rate (V) was measured for each effector in presence of different concentration ([S]) of a fluorescent DNA reporter. Linear fitting of the obtained data revealed that the 2E mutation reduced the *Km* value of the LdCsm-dCsm3 system by 44% (558.44 nM to 313.84 nM) whereas E63A mutation reduces that by 3.2-fold (558.44 nM to 175.14 nM) (Fig. 4, Table 1). These results indicated that reducing negatively charged amino acids in the loops facilitates the substrate binding of the effector, thereby increasing its sensitivity in responding to invading signals.

**Fig. 4.**
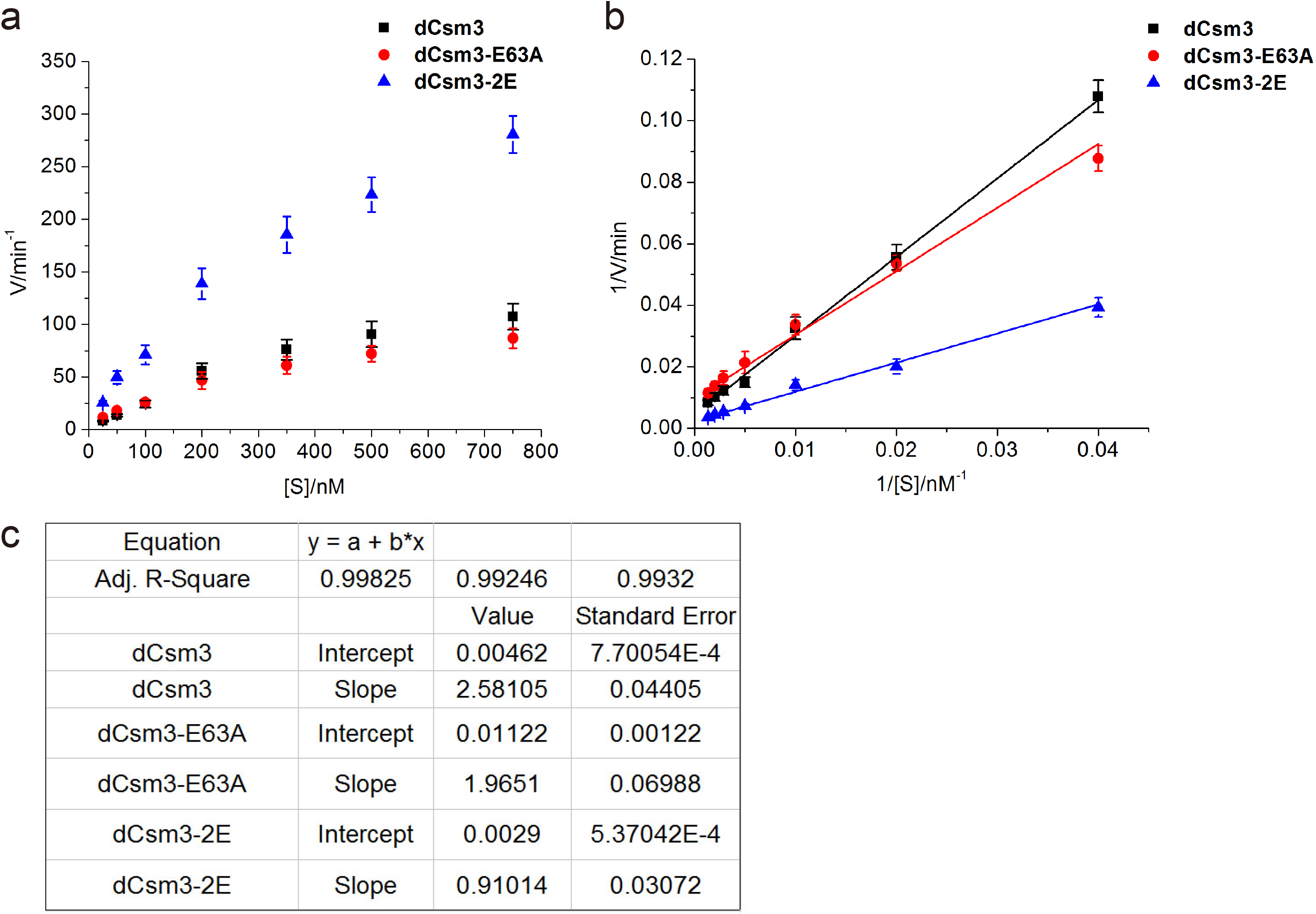
Negative charges in the HD loop segments interfere with ssDNA substrate binding by LdCsm effectors. **a** ssDNA cleavage rates (V) of 50nM LdCsm-dCsm3 (dCsm3), LdCsm-dCsm3-E63A (dCsm3-E63A) or LdCsm-dCsm3-2E (dCsm3-2E) activated by 50nM CTR in presence of different concentrations of fluorescent quenched ssDNA reporter ([S]). **b** Linear fitting of 1/[S] and 1/V. **c** Parameters of linear fitting results. Among the fitting results, value of slope represents *Km*/*Vmax* and value of intercept represents 1/*Vmax*, which can be used to calculate *Km* value that reveals ssDNA substrate binding affinity.

**Table 1.** *Km* values for ssDNA binding determined by kinetic assay.

| LdCsm mutants | dCsm3 | dCsm3-E63A | dCsm3-2E |
| --- | --- | --- | --- |
| *K<sub>m</sub>* value/nM | 558.44 | 175.14 | 313.84 |

dCsm3: LdCsm-dCsm3; dCsm3-E63A: LdCsm-dCsm3-E63A; dCsm3-2E: LdCsm-dCsm3-2E.

### Three HD subdomains contribute to substrate binding

As kinetics assay relies on the activity of a reaction for determination of substrate affinity, the method is only useful for studying active enzymes. Here, we have to evaluate substrate binding activity regardless of whether an LdCsm effector is active or not. Thus, an activity-independent method is required to evaluate the substrate affinity, and MST (Micro-Scale Thermophoresis) assay ^37^ provides such an approach. The method was then employed to test binding of ssDNA substrate by three forms of the WT LdCsm effector, the apo CTR and NTR forms. We found that the binary form (apo-LdCsm) is inactive. Furthermore, the CTR-LdCsm ternary complex binds the substrate with the dissociation constant (*K*_*d*_) value of 2.51 μM whereas NTR-LdCsm interacts with ssDNA in a similar fashion as the apo-form (Table 2, Fig. S6a). Thus, only CTR can activate the substrate binding of LdCsm, and this suggests that substrate binding and cleavage could be a coupled process for LdCsm DNase.

**Table 2.** *Kd* values for ssDNA binding determined by MST assay.

| LdCsm | State | *K<sub>d</sub>*/μM | Signal/Noise |
| --- | --- | --- | --- |
| WT | Apo | ND | - |
| WT | NTR-bound | ND | - |
| WT | CTR-bound | 2.51 ± 2.83 | 7.9 |
| E63A | CTR-bound | 0.93 ± 0.14 | 24.2 |
| K65A | CTR-bound | ND | - |
| K66A | CTR-bound | 2.51 ± 0.98 | 26.3 |
| D94A | CTR-bound | ND | - |
| R95AR96A | CTR-bound | ND | - |
| 2E | CTR-bound | 0.21 ± 0.04 | 6.6 |
| D113N | CTR-bound | ND | - |
| d-L1b | CTR-bound | ND | - |
| d-L1c | CTR-bound | ND | - |
| d-L2c | CTR-bound | ND | - |
| dCsm3-HN | CTR-bound | ND | - |
| dCsm3-HN-3E | CTR-bound | ND | - |

ND: None Detected

Then, MST was employed to re-examine the substrate binding of the mutated effectors showing an increased in vivo interference activity, i.e., E63A and 2E (Fig. S6b). We found that the *K*_*d*_ values of E63A and 2E effectors were ca. 2.7- and 10.2-fold lower than WT, respectively (Table 2). Noticeably, the increase in substrate affinity for the two mutated effectors is now positively related to their ssDNA cleavage activity (CF Fig. 2c∼d and Table 2).

Next, the MST assay was employed to examine substrate binding of LdCsm effectors either with an inactive DNase or a strongly impaired one, including d-L1b, d-L1c, d-L2b, K65A, D94A, R95AR96A and D113N. We found their fitted fluorescence curves in the CTR form are very similar to those obtained with the apo- and NTR-form of WT LdCsm (Fig. S6b). These results indicated that all these mutations strongly impair or completely abolish the substrate binding of LdCsm. To this end, we conclude that the influence of HD loop segments on substrate binding is multifaceted, including the size and amino acid composition of the loops and their remodeling.

The deduced catalytic center for ssDNA cleavage is the HD motif present in Cas10, the largest subunit in type III CRISPR-Cas systems. To investigate if the HD motif could also be involved in substrate binding, we constructed two LdCsm mutants carrying D16N substitution in the HD motif, i.e., dCsm3-Csm1HN (dCsm3-HN) and dCsm3-Csm1HN+E63AE98AE99A (dCsm3-HN-3E). MST analysis of both mutated effectors dCsm3-HN and dCsm3-HN-3E mutants showed no ssDNA binding ability, suggesting that HD catalytic residues are also crucially important to ssDNA binding in LdCsm1 HD domain (Fig. S6b).

Taken together, we demonstrate that the LdCsm DNase is primarily controlled at the stage of substrate binding, which is a complex process involving three subdomains in the HD domain of LdCsm, including HD motif, HD-L1 and HD-L2. In fact, the three subdomains form an extended tripartite substrate-binding motif, and CTR and NTR differentially remodel the tripartite motif to yield activation and inhibition of the LdCsm DNase, respectively.

## Discussion

How target RNA (CTR and NTR) regulates CRISPR-Cas10 DNase has been a long-standing question in the research field. During our investigation on activation mechanisms of the LdCsm DNase, we found that three subdomains are involved in ssDNA substrate binding in the LdCsm system, including HD-L1 and HD-L2, the two flexible loop segments, and the HD catalytic center. We have further revealed that these subdomains work together to capture ssDNA for cleavage and disruption of any of them abolishes substrate binding, yielding a DNase-dead mutant. This extended motif is termed “a tripartite DNA substrate-binding motif”, and it is characterized by the flexibility of two HD loop segments as well as their variable sizes and amino acid sequences. These features provide the molecular basis not only for illustrating the mechanism involved in ssDNA cleavage but also for diversification of the HD domains in CRISPR-Cas10 systems.

Our results, together with the structural features of the HD loops revealed for StCsm ^34^ allow us to propose, for the first time, the molecular mechanisms for CTR activation and NTR inhibition of the LdCsm DNase. In the presence of CTR, the size of the flexible segment is reduced for HD-L1, and in fact, K65, a key amino acid in the partial binding motif of HD-L1, could now interact with DNA substrate from a relatively fixed point, thus increasing the probability of forming two-point bound substrate (Fig. 5a). By contrast, binding of NTR to LdCsm renders HD-L1 even more flexible, and this strongly reduces the propensity for HD-L1 and HD-L2 to bind cooperatively with DNA substrate (Fig. 5a). Thus, remodeling the HD-L1 subdomain by CTR and NTR provides the mechanisms both for CTR activation and for NTR inhibition.

**Fig. 5.**
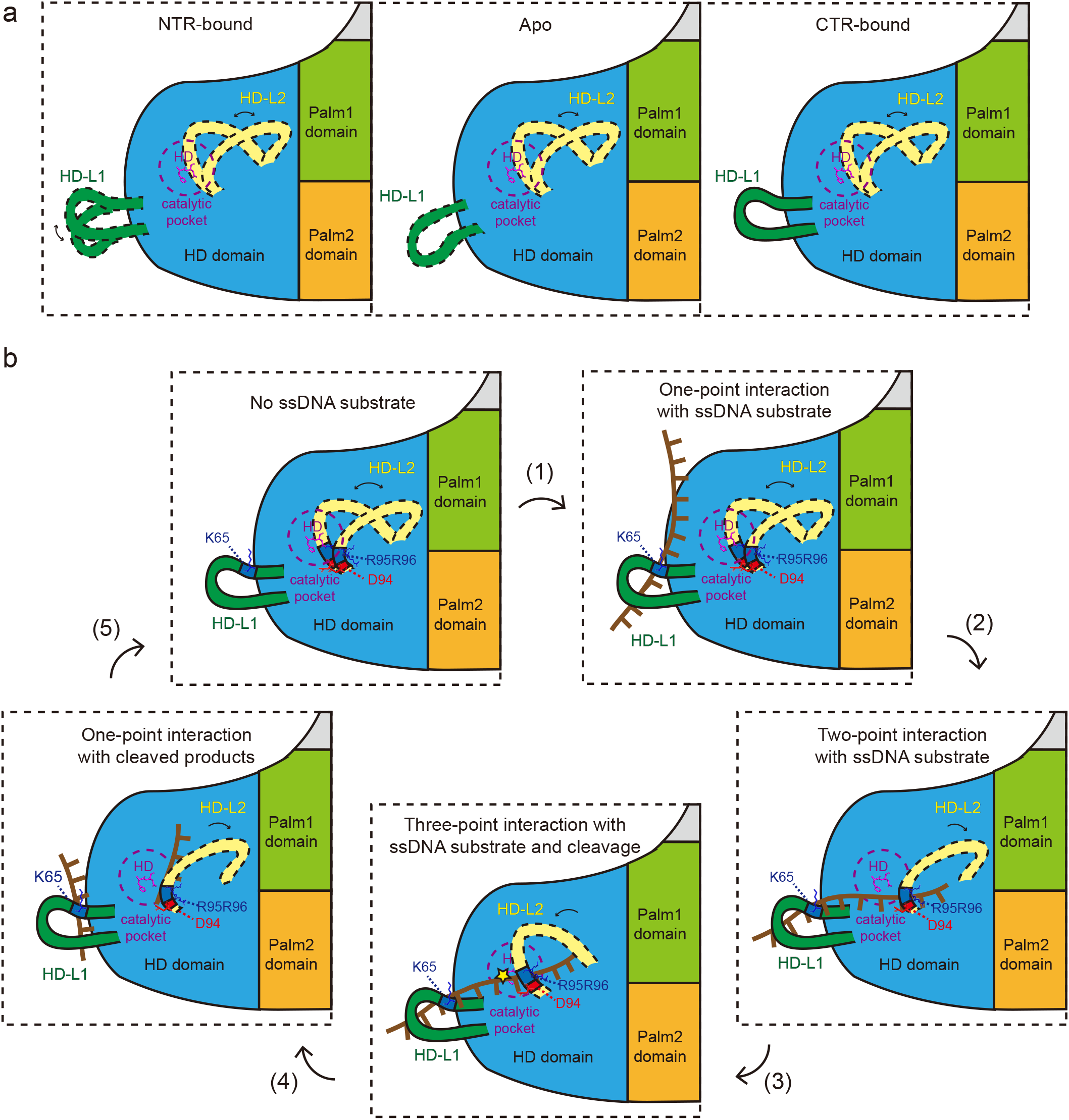
Working model for the mechanism of DNA cleavage by the LdCsm antiviral system. **a** Remodeling of HD-L1 provides the mechanisms for CTR activation and for NTR inhibition of the LdCsm immunization. **b** Deduced steps for ssDNA cleavage in the tripartite substrate-binding motif in the LdCsm HD domain. See detailed explanations in the main text.

The architecture of HD-L2 is very different from HD-L1 since the anchors of the former loop are conserved. One is associated with the GxDRR motif while other is characterized with polar interactions between charged amino acids in the clade of StCsm and LdCsm (Fig. 2c, Fig. S7). These anchors firmly hold the very flexible loop segment of HD-L2 and therefore can support the loop oscillation in a defined pattern. Thus, we infer that HD-L2 could function as a propeller to deliver substrate into the catalytic center and such an action is highly desired, too, for any biochemical reaction to occur in a highly flexible active center of an enzyme.

It is very striking that D16N substitution in the HD motif also completely abolishes the substrate binding capability. This suggests all three subdomains of the LdCsm DNase are involved in substrate binding, including HD-L1, HD-L2 and the HD motif. This provides the basis for the occurrence of an extended tripartite motif, functioning in substrate binding. Indeed, the three subdomains form a triangle in the simulated structures of LdCsm and mutation of any of the binding sites inactivates the DNase, and these findings highlight the importance of cooperation of the three binding sites in the extended tripartite motif.

Taken altogether, we have deduced key steps of DNA cleavage by LdCsm as shown Fig. 5b: (a) one-point substrate-binding is highly dynamic between ssDNA and a binding site in the tripartite motif of LdCsm; (b) an effective reaction starts with the binding of substrate at two loop sites simultaneously; (c) the bound substrate is then propelled into the catalytic site by regular flipping of the HD-L2 loop segment; (d) once the bound substrate is propelled into the catalytic site by HD-L2 flipping, the third binding site in the HD motif interacts with the substrate to yield three point-bound substrate that is relatively stable and ready for cleavage; (e) after cleavage, the resulting cleavage products are either in one-point or in two-point binding forms, which will be readily dissociated from the tripartite motif, thereby completing the DNA cleavage process.

Amongst the structures determined for effector complexes of 4 different III-A CRISPR-Cas10 systems ^34-36, 38^, ToCsm is the only one with a single flexible loop in the HD domain. The HD loop has been captured in the apo or NTR form but is absent from the corresponding CTR form (Fig. S1, Fig. S2b). Thus, it appears that the ToCsm domain organization is different from what we have found with LdCsm. To shot a light on how the tripartite motif model could be applied to other III-A systems, a phylogenetic tree was constructed with a selected set of 140 Csm1 proteins. The analysis has revealed most of these Csm1 proteins (the major group) resemble the architecture of the HD domain of StCsm and LlCsm, whereas only a few others are more closely related to ToCsm (the minor group) (Fig. S7). Analysis of their HD loop segments has revealed that members of the major group carry the GxDRR or xxDRR motif in the predicted HD-L2 (Fig. S2a) whereas those in the minor group possess the consensus of EREE/ERE motif. One of the major differences between the two classes is the HD-L1 fragment is largely fixed in ToCsm (Fig S2b). Since the HD-L1 flexibility is very important for the specificity of CTR activation, the trade-off for its loss in ToCsm would be a poor specificity of the CTR activation, which is indeed observed for mutants of the ToCsm HD domain ^35^. To this end, the architecture of the ToCsm HD domain could be regarded as a simplified version of the identified tripartite motif in which HD-L1 could have largely lost the flexibility during evolution.

Based on the discovery of the tripartite substrate binding motif, we infer that ssDNA substrate should be actively transported into the catalytic site for cleavage. It is thus very interesting to investigate whether such a mechanism could occur for other CRISPR-Cas nucleases. Basically, any occurrence of loop segment in a close proximity of the active center of regulated CRISPR-Cas nucleases could suggest a similar mechanism of active substrate delivery. Thus, conducting such a survey would yield important insights into how nature has invented the strategy of active substrate transport for different enzymes or enzyme complexes.

## Materials and Methods

### Bacterial strains and growth conditions

*E. coli* DH5α was the bacterial host for DNA cloning. *E. coli* BL21(DE3) was the host for conducting interference plasmid assay and for expression and purification of LdCsm effector complexes. *E. coli* strains were cultured in Luria-Bertani (LB) or terrific broth (TB) medium. Incubation was at 37 °C, 200 rpm. If required, antibiotics were added as the following: ampicillin (Amp) at 100 μg/ml, kanamycin (Kan) at 25 μg/ml, and chloramphenicol (Cm) at 10 μg/ml.

### Construction of plasmids for the in vivo LdCsm genetic assay

Four new plasmids were constructed for in vivo genetic assay in this work, include LdCsm-expressing plasmid pColE-Cas-S1, and three target/reference plasmids, including p15A-CTR: cognate target RNA-expressing plasmid, p15A-NTR: noncognate target RNA-expressing plasmid, and p15A-G: a reference plasmid that does not carry any target sequence. Their constructions were based on p15A-Cas-S1 and pColE-G/CTR/NTR plasmids reported previously ^31^.

In the construction of pColE-Cas-S1, a ColE backbone fragment (ColE1 origin plus a kanamycin-resistant gene) was amplified from pColE-CTR by PCR using primers ColE-F and ColE-R; the LdCsm-expressing cassette was obtained from p15A-Cas-S1 by PCR with the primer set of Cas-F and Cas-R in pColE-Cas-S1. The two fragments were assembled together via 18∼20nt homologous sequences at their ends using a Gibson assembly system (ABclonal Technology, Wuhan, China), giving pColE-Cas-S1.

For construction of p15A series plasmids, a backbone fragment (p15A origin plus an ampicillin-resistant marker) was obtained from p15A-Cas-S1 by PCR, using primers p15A-F and p15A-R. The insert fragments were amplified from pColE-G/CTR/NTR ^31^, respectively, using the primer set of arab-F and arab-R (sequences of primers are listed in Table S3). Then, Gilson assemble was employed for generation of these plasmids, yielding p15A-G/CTR/NTR.

### Plasmid interference assay

*Escherichia coli* BL21(DE3) strain carrying pColE-Cas-S1 and p15A-CTR was named as test strain (T2) to reveal the in vivo interference activity of LdCsm and its mutated derivatives. Another strain containing pColE-Cas-S1 and p15A-G (lacking any target sequence) was used as a reference (CK2). A third strain carrying pColE-Cas-S1 and p15A-NTR was for testing the influence of noncognate target RNA on the immunity. Single colonies of test strain and reference strains were inoculated into 5 ml LB medium containing 100μg/ml Amp and 50μg/ml Kan and cultured overnight at 37 °C 220 rpm. Then, the seed cultures were transferred to 30 ml LB medium without any selection and cultured to mid-log phase (OD_600_=0.8) at 37 °C 220 rpm. L-arabinose was then added to 0.1% to induce the transcription of CTR which would then activate the LdCsm immunization. Samples were taken every 0.5h/0.75h, which were then plated onto LB agar plates without antibiotics, or in the presence of 100μg/ml Amp, or 50μg/ml Kan. After incubation at 37 °C overnight, colonies were enumerated with which the survival rates were calculated for each strain.

### Construction of *cas* gene plasmids carrying mutations in the HD domain of the *L. delbrueckii* Csm1

Eight HD loop truncations and 14 substitutions were designed for the Csm1 HD domain, including d-L1a, d-L1b, d-L1c, d-L1d, d-L2a, d-L2b, d-L2c, d-L2d, E63A, K65A, K66A, K65AK66A, R70A, E73AE76A, D94A, R95AR96A, E98AE99A, K102A, D106A, D113A, D113N or R125A mutation in the Csm1-encoding sequence. They all were constructed using a PCR and Gibson assembly approach that was briefly described below. Two sets of complementary primers were designed: one was at the mutation position (for a variant *csm1* gene with mutation X, the two primers were named X-F and X-R), the other set of complementary primers was picked up from the Csm5 coding sequence (called over-F and over-R) (sequences of primers are listed in Table S3). Then, a smaller fragment of pColE-Cas-S1 was amplified from pColE-Cas-S1 by PCR using the X-F and over-R primers while a larger fragment was amplified using over-F and X-R from the same plasmid. Ligation of two fragments by Gibson assembly yielded pColE-Cas-S1 carrying mutation X. In addition, using p15AIE-Cas as the template for the PCR reactions produced the p15AIE-Cas plasmids carrying each mutation designed for the Csm1 protein. The two series of plasmids were to be used in genetic assays and for RNP purification, respectively.

### Purification of LdCsm effector complexes from *E. coli*

LdCsm purification was conducted as previously described ^31^. Expression strains were obtained by transformation of *E. coli* BL21/DE3 with three plasmids (p15AIE-Cas or its derivative, pET30a-Csm2 and pUCE-S1). Several single colonies were picked up from the selective plates and inoculated into 20 ml LB medium with antibiotics. Cultivation was at 37 °C, 220 rpm overnight. 10 ml of the overnight culture was transferred to 1 L TB medium and cultured to a mid-log phase (OD_600_=0.8) under the same growth condition. Then, IPTG was added to 0.3 mM and the culture was further cultured at 25 °C for 16 h to induce the synthesis of LdCsm. Cells were harvested by centrifugation at 5000 rpm for 5 min, and cell pellets were resuspended in 50 ml buffer A (20 mM Tris-HCl, 0.25 M NaCl, 20 mM imidazole, 10% glycerol, pH8.5). The resulting cell suspension was treated by French press at 4°C and cell debris was removed by centrifugation at 10,000 rpm for 1 h at 4 °C. LdCsm complexes were purified in a two-step purification procedure as previously described^31^.

The LdCsm complex is captured on the HisTrap affinity column and eluted with buffer B (20 mM Tris-HCl, 0.25 M NaCl, 200 mM imidazole, 10% glycerol, pH8.5). The product is further purified by gel filtration with Superdex 200 (GE Healthcare, Little Chalfont, United Kingdom) using the chromatography buffer (20 mM Tris-HCl, 0.25 M NaCl, 5% glycerol, pH8.5). Fraction samples are collected and used for further analysis.

### Nucleic acid cleavage assay

Nucleic acid cleavage assays (10 μl) were conducted as the following described. DNA cleavage reactions were conducted in the cleavage buffer of 50 mM Tris-HCl, 10 mM MgCl2, 50 mM KCl, 0.1 mg/ml BSA, pH7.0 containing the amounts of the WT LdCsm or one of its derivative effectors indicated in each assay, 50 nM 5’ FAM-labeled ssDNA substrate, and 500 nM target RNA (sequences of target RNAs are listed in Table S4). For RNA cleavage assay, indicated amounts of an LdCsm effector were mixed with 50 nM 5’ FAM-labeled target RNA in the cleavage buffer (50 mM Tris-HCl, 10 mM MgCl2, 50 mM KCl, 0.1 mg/ml BSA, pH7.0). Nucleic acids cleavage was conducted at 37 °C for 10 min and stopped by addition of 2×RNA loading dye (New England Biolabs, Ipswich, MA, USA). Before loading, samples were heated for 3 min at 95 °C and analyzed by denaturing PAGE. Cleavage products were visualized by fluorescence imaging analysis of the gel.

### Fluorescence DNA reporter cleavage assay

The DNA reporter assay employed the same reaction setup as the DNA cleavage reaction described above except that 1 μM FAM-poly-16T-BHQ1 ssDNA substrate (Azenta Biotechnology Company, Qingdao, China) was used as DNA substrate and the total reaction volume was 20 μl. Reporter reactions were set up in a 384-well black plate (Thermo fisher, Waltham, MA, USA) and incubated in an Enspire fluorescence plate reader (Perkin Elmer, Waltham, MA, USA) for 30∼60 min at 37 °C. Fluorescence was measured in every 5/10 min (λ_ex_: 485 nm; λ_em_: 535 nm). Relative activity was obtained by subtraction of the background fluorescence values generated for the reference sample without any target RNA from the fluorescence values obtained from the reactions containing the cognate target RNA or RNAs to be detected.

### Micro-scale thermophoresis assay of LdCsm substrate binding

MST experiments were carried out at 22 °C on Monolith NT.115 instrument (NanoTemper Technologies, Munich, Germany) as described in the review entitled *Molecular interaction studies using microscale thermophoresis* ^37^. The concentration of wild type LdCsm or mutants with equal amount of cognate target RNA (CTR) were gradient-diluted from 4μM to 0.122 nM, while a 30nt 5’-FAM labelled ssDNA substrate was added to constant concentration of 5 nM. After a short incubation in buffer containing 50 mM Tris-HCl (pH 7.0), 125 mM NaCl and 25mM Ethylenediamine tetraacetic acid (EDTA)The sample was loaded into the Monolith NT.115 Capillary. Micro thermophoresis was carried out using 100% LED power and 40% MST power. Dissociation constant *Kd* of the wild type LdCsm and its mutated derivatives were calculated using the MO software (MO. Affinity Analysis (x86), NanoTemper Technologies, Munich, Germany) and data fitting was performed using OriginPro 2015 software (OriginLab, Northampton, MA, USA).

### Phylogenetic relationship analysis of type III-A Csm1

Homologs of LdCsm1 were found by protein BLAST (Basic Local Alignment Search Tool) (https://blast.ncbi.nlm.nih.gov/Blast.cgi) in NCBI (National center for biotechnology information) website. Protein sequences of 140 Csm1 from type III-A systems in different species were downloaded (information of the 140 Csm1 were placed in Table S5). These sequences were compared multiplexed by the ClustalW matching method in MEGA7.0 ^39^. Then the result was used to construct the phylogenetic tree using the Maximum Likelihood method with parameters as default.

### Statistical analysis

In vivo assays and biochemical assays were repeated three times, and one representative data are shown.

## Materials availability

Plasmids generated in this study are stored in the CRISPR and Archaea Biology Research Center, State Key Laboratory of Microbial Technology, Shandong University, Qingdao 266237, China. Any requests can be made by contacting the corresponding author.

## Acknowledgements

This study was supported by grants from the National Key R & D Program of China (2021YFA0717000), the National Natural Science Foundation of China (Grant No. 31771380) to QS; from the Chinese Postdoctoral Science Foundation (2020M672050), the Qingdao Applied Research Fund For Postdoctoral Researchers (62450079311107) to ZY, and an Open Project from the State Key Laboratory of Microbial Technology at Shandong University.

## Author Contributions

Z.Y. designed the experiments, carried out most of the experiments, analyzed the data, prepared figures and drafted a first version of the manuscript. F.W. carried out the MST assay and analyzed the data, participated in structure modelling and optimized the illustration of structural models. Q.S. supervised the research, designed the experiments, analyzed the research data, wrote and revised the manuscript with the help from Z.Y. and F.W.

## References

1 Marraffini LA. CRISPR-Cas immunity in prokaryotes. Nature 2015; 526:55–61.

2 Mohanraju P, Makarova KS, Zetsche B, Zhang F, Koonin EV, van der Oost J. Diverse evolutionary roots and mechanistic variations of the CRISPR-Cas systems. Science 2016; 353:aad5147.

3 Wright AV, Nunez JK, Doudna JA. Biology and Applications of CRISPR Systems: Harnessing Nature’s Toolbox for Genome Engineering. Cell 2016; 164:29–44.

4 Hille F, Richter H, Wong SP, Bratovič M, Ressel S, Charpentier E. The Biology of CRISPR-Cas: Backward and Forward. Cell 2018; 172:1239–1259.

5 Makarova KS, Wolf YI, Iranzo J et al. Evolutionary classification of CRISPR-Cas systems: a burst of class 2 and derived variants. Nature reviews Microbiology 2020; 18:67–83.

6 Marraffini LA, Sontheimer EJ. CRISPR interference limits horizontal gene transfer in staphylococci by targeting DNA. Science (New York, NY) 2008; 322:1843–1845.

7 Hale CR, Zhao P, Olson S et al. RNA-guided RNA cleavage by a CRISPR RNA-Cas protein complex. Cell 2009; 139:945–956.

8 Deng L, Garrett RA, Shah SA, Peng X, She Q. A novel interference mechanism by a type IIIB CRISPR-Cmr module in Sulfolobus. Molecular microbiology 2013; 87:1088–1099.

9 Peng W, Feng M, Feng X, Liang YX, She Q. An archaeal CRISPR type III-B system exhibiting distinctive RNA targeting features and mediating dual RNA and DNA interference. Nucleic acids research 2015; 43:406–417.

10 Tamulaitis G, Kazlauskiene M, Manakova E et al. Programmable RNA shredding by the type III-A CRISPR-Cas system of Streptococcus thermophilus. Molecular cell 2014; 56:506–517.

11 Tamulaitis G, Venclovas C, Siksnys V. Type III CRISPR-Cas immunity: major differences brushed aside. Trends in microbiology 2017; 25:49–61.

12 Zhang Y, Lin J, Feng M, She Q. Molecular mechanisms of III-B CRISPR–Cas systems in archaea. Emerging Topics in Life Scienes 2018; 2:483–491.

13 Benda C, Ebert J, Scheltema RA et al. Structural model of a CRISPR RNA-silencing complex reveals the RNA-target cleavage activity in Cmr4. Molecular cell 2014; 56:43–54.

14 Zhu X, Ye K. Cmr4 is the slicer in the RNA-targeting Cmr CRISPR complex. Nucleic acids research 2015; 43:1257–1267.

15 Samai P, Pyenson N, Jiang W, Goldberg GW, Hatoum-Aslan A, Marraffini LA. Co-transcriptional DNA and RNA Cleavage during Type III CRISPR-Cas Immunity. Cell 2015; 161:1164–1174.

16 Elmore JR, Sheppard NF, Ramia N et al. Bipartite recognition of target RNAs activates DNA cleavage by the Type III-B CRISPR-Cas system. Genes & development 2016; 30:447–459.

17 Estrella MA, Kuo FT, Bailey S. RNA-activated DNA cleavage by the Type III-B CRISPR-Cas effector complex. Genes & development 2016; 30:460–470.

18 Kazlauskiene M, Tamulaitis G, Kostiuk G, Venclovas C, Siksnys V. Spatiotemporal Control of Type III-A CRISPR-Cas Immunity: Coupling DNA Degradation with the Target RNA Recognition Molecular cell 2016; 62:295–306.

19 Han W, Li Y, Deng L et al. A type III-B CRISPR-Cas effector complex mediating massive target DNA destruction. Nucleic acids research 2017; 45:1983–1993.

20 Niewoehner O, Garcia-Doval C, Rostol JT et al. Type III CRISPR-Cas systems produce cyclic oligoadenylate second messengers. Nature 2017; 548:542–548.

21 Kazlauskiene M, Kostiuk G, Venclovas C, Tamulaitis G, Siksnys V. A cyclic oligonucleotide signaling pathway in type III CRISPR-Cas systems. Science 2017; 357:605–609.

22 Han W, Stella S, Zhang Y et al. A Type III-B Cmr effector complex catalyzes the synthesis of cyclic oligoadenylate second messengers by cooperative substrate binding. Nucleic acids research 2018; 46:10319–10330.

23 Rouillon C, Athukoralage JS, Graham S, Gruschow S, White MF. Control of cyclic oligoadenylate synthesis in a type III CRISPR system. Elife 2018; 7:e36734.

24 Grüschow S, Athukoralage JS, Graham S, Hoogeboom T, White MF. Cyclic oligoadenylate signalling mediates Mycobacterium tuberculosis CRISPR defence. Nucleic acids research 2019; 47:9259–9270.

25 Rostol JT, Marraffini LA. Non-specific degradation of transcripts promotes plasmid clearance during type III-A CRISPR-Cas immunity. Nature microbiology 2019; 4:656–662.

26 Lau RK, Ye Q, Birkholz EA et al. Structure and Mechanism of a Cyclic Trinucleotide-Activated Bacterial Endonuclease Mediating Bacteriophage Immunity. Molecular cell 2020; 77:723-733.e726.

27 McMahon SA, Zhu W, Graham S, Rambo R, White MF, Gloster TM. Structure and mechanism of a Type III CRISPR defence DNA nuclease activated by cyclic oligoadenylate. Nature communications 2020; 11:500.

28 Rostol JT, Xie W, Kuryavyi V et al. The Card1 nuclease provides defence during type III CRISPR immunity. Nature 2021; 590:624–629.

29 Zhu W, McQuarrie S, Gruschow S et al. The CRISPR ancillary effector Can2 is a dual-specificity nuclease potentiating type III CRISPR defence. Nucleic acids research 2021.

30 Molina R, Stella S, Feng M et al. Structure of Csx1-cOA4 complex reveals the basis of RNA decay in Type III-B CRISPR-Cas. Nature communications 2019; 10:4302.

31 Lin J, Feng M, Zhang H, She Q. Characterization of a novel type III CRISPR-Cas effector provides new insights into the allosteric activation and suppression of the Cas10 DNase. Cell Discov 2020; 6:29.

32 Lin J, Shen Y, Ni J, She Q. A type III-A CRISPR-Cas system mediates co-transcriptional DNA cleavage at the transcriptional bubbles in close proximity to active effectors. Nucleic acids research 2021; 49:7628–7643.

33 Molina R, Sofos N, Montoya G. Structural basis of CRISPR-Cas Type III prokaryotic defence systems. Current opinion in structural biology 2020; 65:119–129.

34 You L, Ma J, Wang J et al. Structure Studies of the CRISPR-Csm Complex Reveal Mechanism of Co-transcriptional Interference. Cell 2019; 176:239-253.e216.

35 Jia N, Mo CY, Wang C, Eng ET, Marraffini LA, Patel DJ. Type III-A CRISPR-Cas Csm Complexes: Assembly, Periodic RNA Cleavage, DNase Activity Regulation, and Autoimmunity. Molecular cell 2019; 73:264-277.e265.

36 Sridhara S, Rai J, Whyms C et al. Structural and biochemical characterization of in vivo assembled Lactococcus lactis CRISPR-Csm complex. Communications biology 2022; 5:279.

37 Jerabek-Willemsen M, Wienken CJ, Braun D, Baaske P, Duhr S. Molecular interaction studies using microscale thermophoresis. Assay and drug development technologies 2011; 9:342–353.

38 Sofos N, Feng M, Stella S et al. Structures of the Cmr-beta complex reveal the regulation of the immunity mechanism of type III-B CRISPR-Cas. Molecular cell 2020; 79:741–757 e747.

39 Kumar S, Stecher G, Tamura K. MEGA7: Molecular Evolutionary Genetics Analysis Version 7.0 for Bigger Datasets. Molecular biology and evolution 2016; 33:1870–1874.

40 Thompson JD, Higgins DG, Gibson TJ. CLUSTAL W: improving the sensitivity of progressive multiple sequence alignment through sequence weighting, position-specific gap penalties and weight matrix choice. Nucleic acids research 1994; 22:4673–4680.

41 Gouet P, Robert X, Courcelle E. ESPript/ENDscript: Extracting and rendering sequence and 3D information from atomic structures of proteins. Nucleic acids research 2003; 31:3320–3323.

