## Supplementary information for "Remodeling of a tripartite substrate-binding motif in the HD domain provides the mechanism for activation of CRISPR-Cas10 DNases"

**Description of supplementary information:**

Seven supplementary figures (Fig. S1~Fig. S7);

Five supplementary tables (Table S1~Table S5).

### Supplementary figures

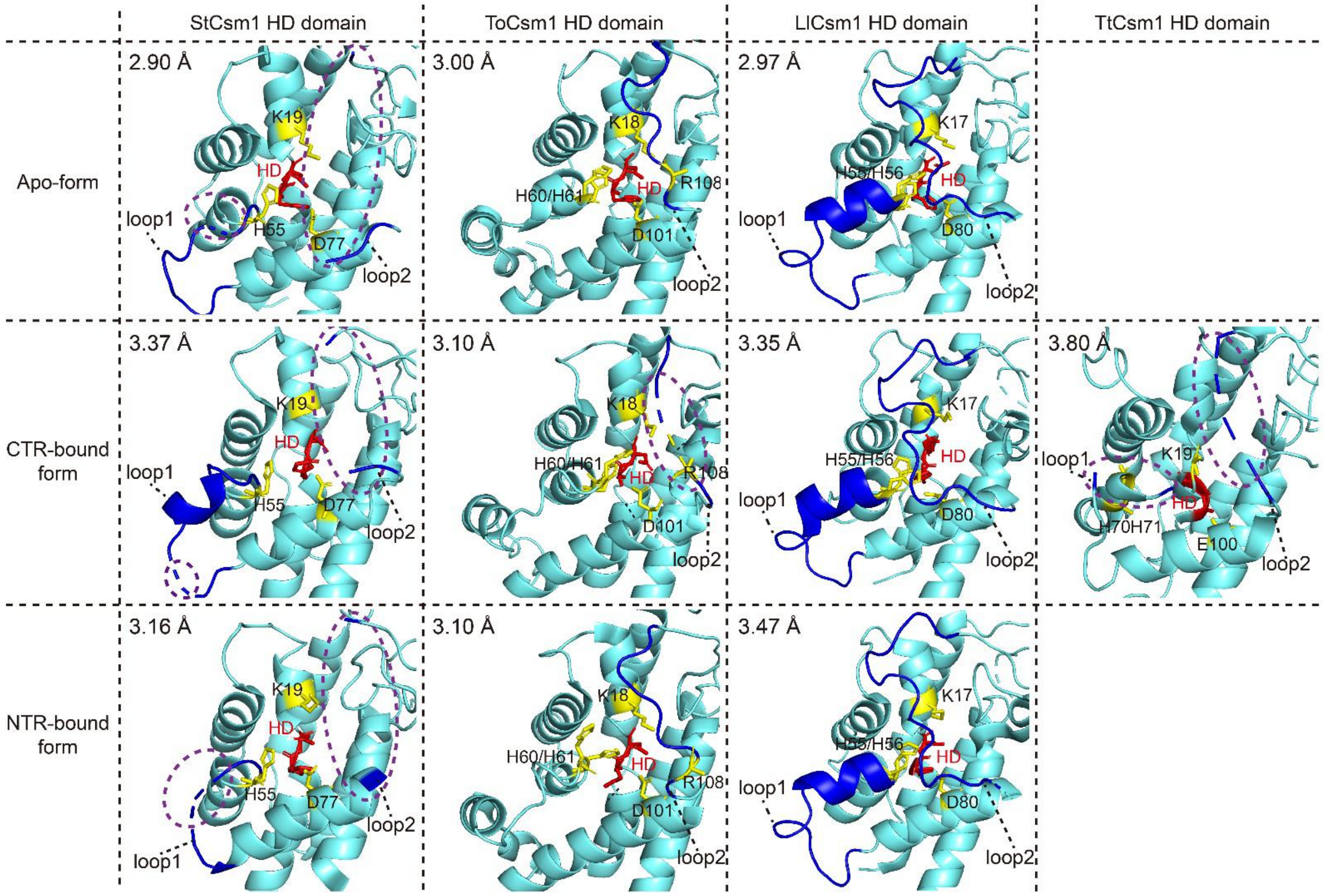


**Fig. S1 Structural comparisons of loop segments present in the Csm1 HD domain of four known Csm structures. Related to Fig. 1**

Structure data of StCsm (PDB ID: 6ifn, 6ify, 6ifl), ToCsm (PDB ID: 6muu, 6mur, 6mut), LlCsm (PDB ID: 6xn5, 6xn3, 6xn7) and TtCsm (PDB ID: 6o1o) were downloaded from PDB database and viewed with the PyMol program. Only the HD domain regions are shown. Resolution of the structures are labeled in the upper left corner in each panel. Active sites of the HD domain are highlighted in red; indispensable amino acid residues identified for the DNase catalytic pocket of ToCsm1 are highlighted in yellow, along with their corresponding amino acid residues in StCsm1, LlCsm1 and TtCsm1. Loop structures present in the HD domains are highlighted in blue. Loop segments cannot be traced are indicated with purple dash line circle.

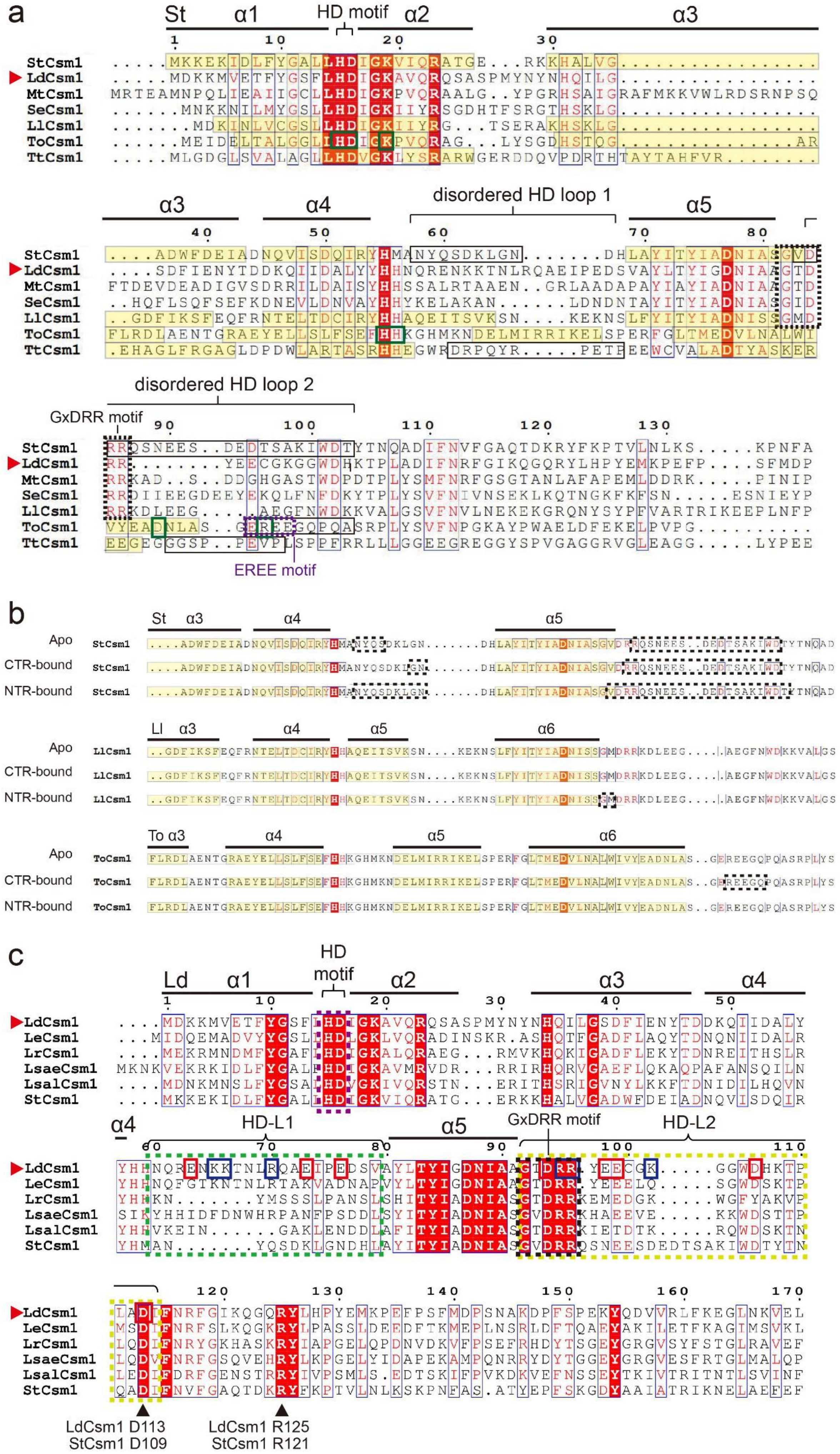
a
St α1 HD motif α2 α3
StCsm1
LdCsm1
MtCsm1
SeCsm1
LlCsm1
ToCsm1
TtCsm1
α3 α4 disordered HD loop 1 α5
disordered HD loop 2
GxDRR motif
EREE motif
b
St α3 α4 α5
Apo
CTR-bound
NTR-bound
Ll α3 α4 α5 α6
To α3 α4 α5 α6
c
Ld α1 HD motif α2 α3 α4
LdCsm1
LeCsm1
LrCsm1
LsaeCsm1
LsalCsm1
StCsm1
α4 HD-L1 α5 GxDRR motif HD-L2
LdCsm1 D113
StCsm1 D109
LdCsm1 R125
StCsm1 R121

**Fig. S2 Conservation and diversification of HD loop segments in different type III-A systems. Related to Fig. 1**

**a** Structure-based sequence alignment of a selected set of Csm1 proteins. Flexible disordered loops are boxed in solid lines; GxDRR motifs at the start position of HD loop 2 are boxed in dash lines. EREE motif at the start position of the single loop in ToCsm1 is boxed in purple dash lines. Amino acid residues characterized for the ToCsm1 HD domain are boxed in solid dark green lines. **b** Summary of untraceable HD loop segments (boxed in dash lines) in the apo, CTR- or NTR-Csm complex for StCsm1, LlCsm1 and ToCsm1 (PDB ID of StCsm complexes: 6ifn, 6ify, 6ifl; PDB ID of LlCsm complexes: 6xn5, 6xn3, 6xn7; PDB ID of ToCsm complexes: 6muu, 6mur, 6mut). **c** An alignment of LdCsm1 and homologous Csm1 sequences. Two disordered loops in the HD region (HD-L1, HD-L2) are boxed in light green and dark yellow dashed lines, whereas positively and negatively charged amino acids in the loops are highlighted in blue boxes and red boxes for LdCsm1. Csm1 sequences selected for analysis include those from type III-A systems of *Streptococcus thermophilus* (St), *Lactobacillus delbrueckii* (Ld), *Mycobacterium tuberculosis* (Mt), *Staphylococcus epidermidis* (Se), *Lactococcus lactis* (Ll), *Thermococcus onnurineus* (To), *and Thermus thermophilus* (Tt). *Lactobacillus salivarius* (Lsal), *Lactobacillus equicursoris* (Le), *Lactobacillus ruminis* (Lr), and *Lactobacillus saerimneri* (Lsae). LdCsm1 sequence is marked by red triangle. Sequence alignment was conducted using CLUSTAL W [40] and Endscript [41].

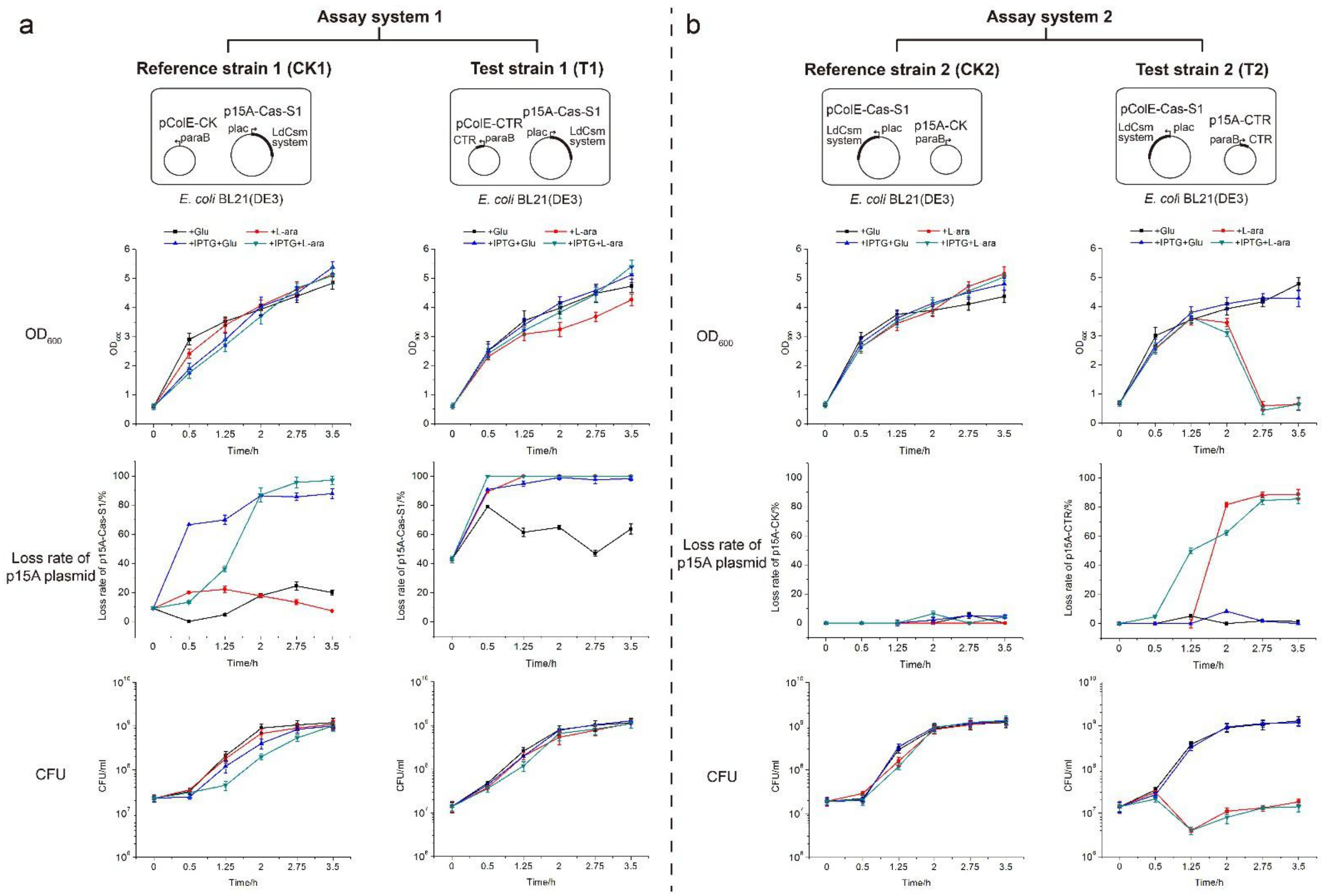


**Fig. S3 Evaluation of LdCsm immune responses using two in vivo assay systems. Related to Fig. 1**

**a** Evaluation of LdCsm immune responses using the reported in vivo assay system (Lin et al. 2021). Test strain in this assay carrying p15A-Cas-S1 and pColE-CTR whereas the corresponding reference strain containing p15A-Cas-S1 and pColE-CK. Samples were taken from cultures of each strain for determination of $OD_{600}$ values, plasmid loss rates and colony forming units (CFU) at different times during incubation. Each strain was cultured to a mid-log phase ($OD_{600}$=0.8) at 37 ºC, 220 rpm in the absence of antibiotics and divided into four equal portions for conducting the experiment. p15A-Cas1-S1: low copy plasmid for expression of the LdCsm system from the *lac* promoter; pColE-CTR: high copy plasmid for expression of cognate target RNA from the *araBad* promoter. **b** Evaluation of LdCsm immune responses using a new plasmid interference assay. Test strain in the new plasmid interference assay carrying pColE-Cas-S1 and p15A-CTR and its reference strain containing pColE-Cas-S1 and p15A-CK. Compared with the reported assay, the only difference is that plasmid backbones for expression of LdCsm effector and CTR were exchanged. L-ara: L-arabinose (0.1%), IPTG: *lac* promoter-inducer added to 0.3mM, Glu: glucose (0.5%) for mediating the catabolite repression to inhibit residual expression from the *lac* and *araBad* promoters.

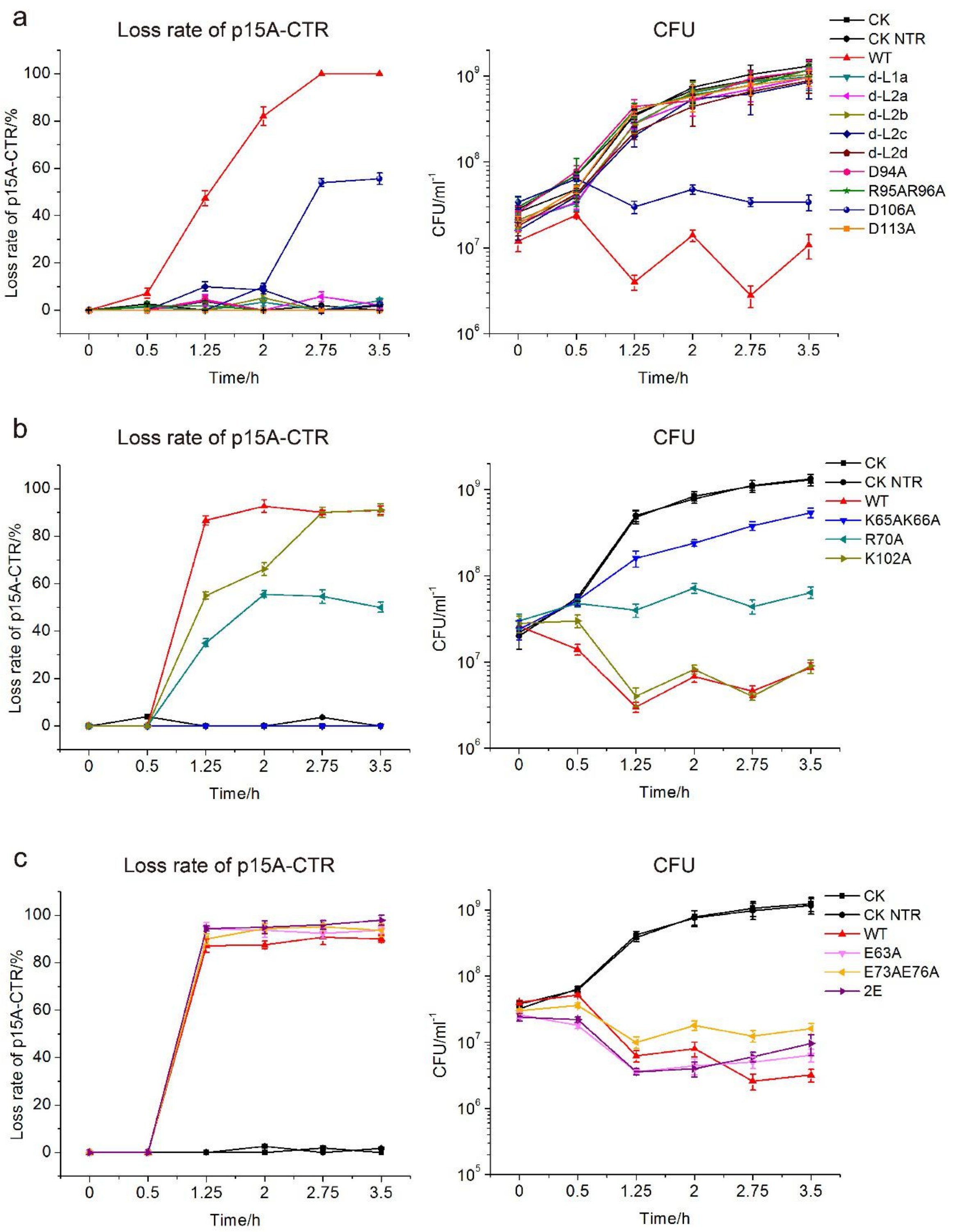


**Fig. S4 Functional analysis of the HD loops using the new LdCsm interference assay. Related to Fig. 1**
**a** Analysis of test strains carrying HD-L1/L2 truncations or mutations on conserved amino acid residues in HD-L2 in the new LdCsm interference assay. **b** Analysis of test strains carrying mutations on positively charged amino acid residues in HD-L1/L2 in the new LdCsm interference assay. **c** Analysis of test strains carrying mutations on negatively charged amino acid residues in HD-L1/L2 in the new LdCsm interference assay. Target plasmid loss rate and colony forming unit (CFU) of test strains at different time points post induction. Wild type test strain (WT) contains pColE-Cas-S1 and p15A-CTR. Mutation X test strain (X) contains pColE-Cas-S1-X and p15A-CTR. Reference strain (CK) contains pColE-Cas-S1 and p15A-CK. NTR reference strain (CK NTR) carries pColE-Cas-S1 and p15A-NTR.

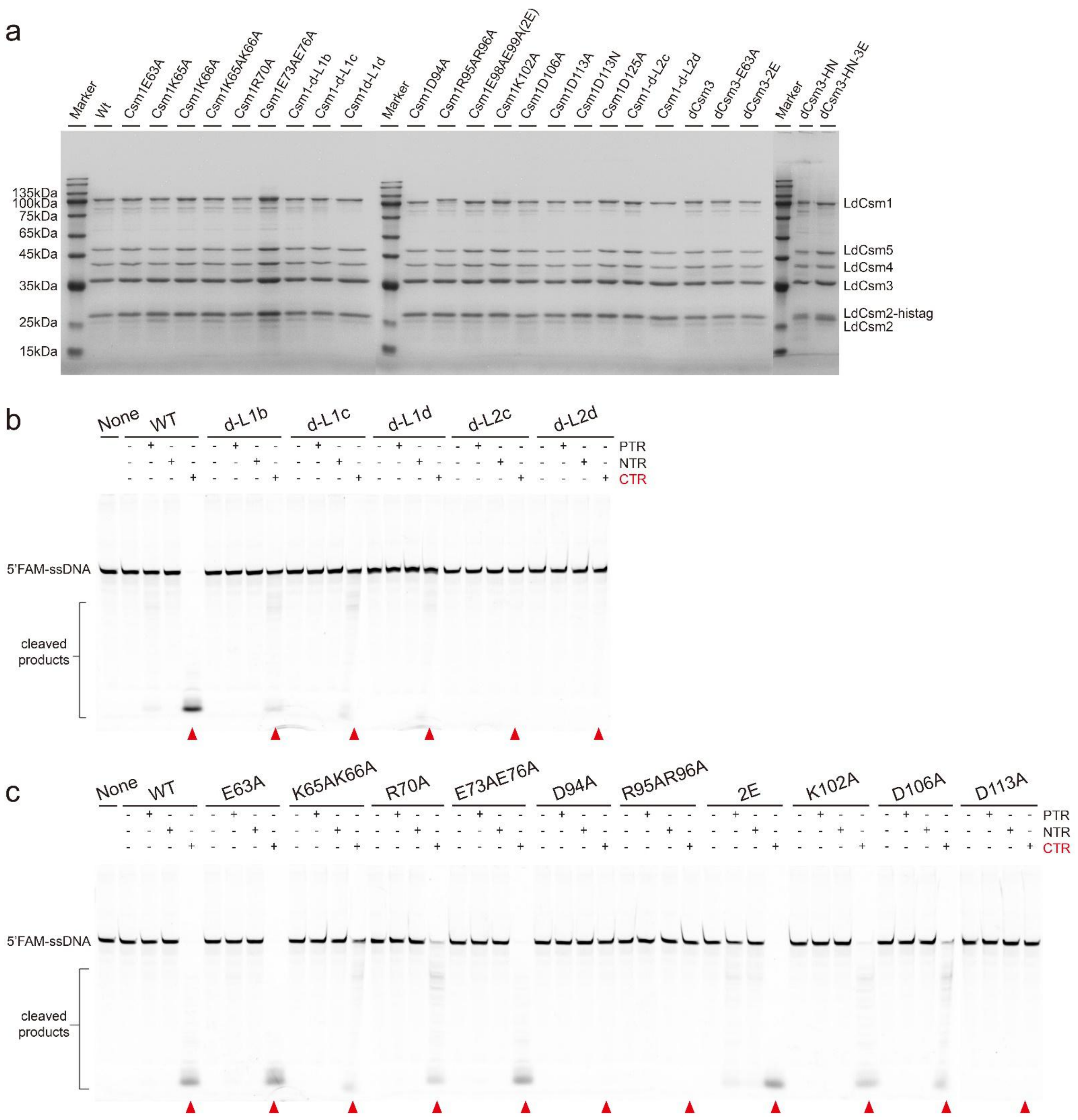


**Fig. S5 DNA cleavage assay of LdCsm mutants carrying truncations of the HD loops or mutations on charged amino acids in the HD loops. Related to Fig. 2**

**a** SDS-PAGE of all the LdCsm mutants purified in this work. The mutations carried by each LdCsm effector is indicated on the top of the gel. dCsm3: nuclease-dead Csm3; dCsm3-E63A: nuclease-dead Csm3-Csm1E63A; dCsm3-2E: nuclease-dead Csm3-Csm1E98AE99A; dCsm3-HN: nuclease-dead Csm3-Csm1D16N (HD domain); dCsm3-HN-3E: nuclease-dead Csm3-Csm1D16N (HD domain) E63AE98AE99A. **b** Target RNA-activated ssDNA cleavage by LdCsm effectors of WT and HD loops truncations. **c** The ssDNA cleavage by LdCsm effectors of WT and amino acid substitutions of HD loops. Cleavage reactions were conducted in 50 nM of LdCsm effector, 50 nM 5' FAM-labeled S10-60 ssDNA substrate and 500 nM PTR, CTR or NTR, and incubation was for 20~30 min at 37$^{o}$C. Cleavage products were analyzed by denaturing PAGE. Red triangles: DNA cleavage reactions activated by CTR, which is also highlighted in red.

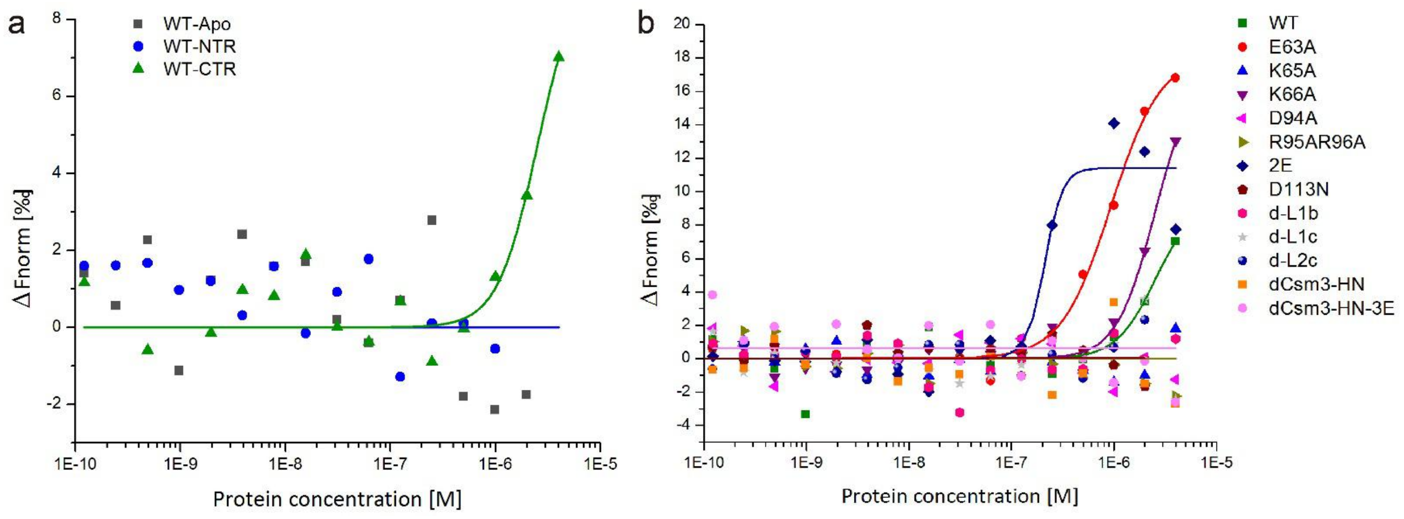


**Fig. S6 MST analysis of interaction between ssDNA substrates and the WT LdCsm or each of the listed LdCsm mutants. Related to Table 2**

**a** MST data of ssDNA substrate binding by Apo, NTR-bound or CTR-bound form of the WT LdCsm. **b** MST data of ssDNA substrate binding by CTR-WT LdCsm and each of CTR-mutated LdCsm effectors. Experiments were repeated three times and showed consistent results and representative data fitting was performed using OriginPro 2015 software. Mutations carried in each LdCsm mutant are annotated and highlighted with different shapes and colors.

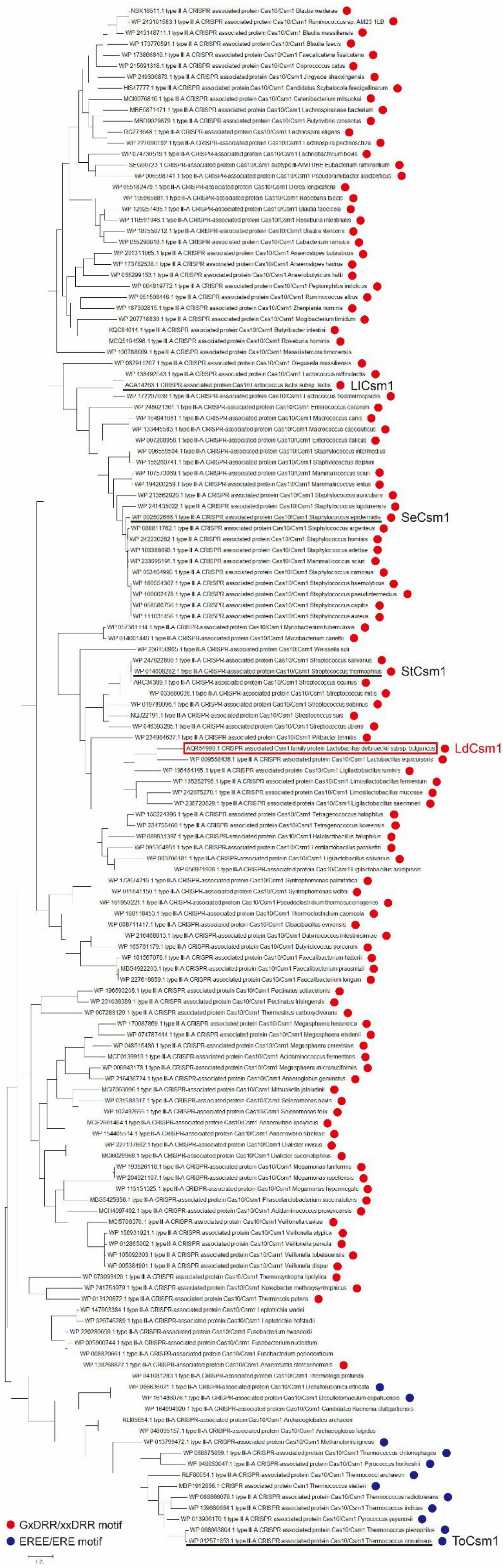

LlCsm1
SeCsm1
StCsm1
LdCsm1
ToCsm1
GxDRR/xxDRR motif
EREE/ERE motif

**Fig. S7 Molecular phylogenetic analysis of type III-A Csm1. Related to Table S5**

LdCsm1 is boxed by red line. StCsm1, SeCsm1, LlCsm1 and ToCsm1 are underlined by black line. Red dots indicate Csm1 homologs carrying GxDRR/xxDRR motif in HD-L2. Blue dots indicate Csm1 homologs carrying EREE/ERE motif in HD-L2. Information of Csm1 used in this figure are listed in Table S5. Molecular phylogenetic analysis of Csm1 was performed by MEGA7.0 [39] using Maximum Likelihood method.

**Supplementary tables**

**Table S1. Parameters of structural modeling of LdCsm1 using different characterized Csm1 as templates. Related to Fig. 1.**

| Template | Seq Similarity | Coverage | GMQE | QMEANDisCo Global |
|---|---|---|---|---|
| SthCsm1 | 0.4 | 0.93 | 0.68 | 0.66 ± 0.05 |
| ToCsm1 | 0.32 | 0.85 | 0.54 | 0.58 ± 0.05 |
| LlCsm1 | 0.37 | 0.93 | 0.63 | 0.61 ± 0.05 |

Structural modeling of LdCsm1 was performed by SWISS-MODEL (https://swissmodel.expasy.org/) (Waterhouse et al., 2018) using different templates (PDB files 6ify (StCsm), 6umr (ToCsm), 6xn4 (LlCsm)). GMQE and QMEANDisCo global give an overall model quality measurement between 0 and 1, with higher numbers indicating higher expected quality. GMQE is coverage dependent, i.e., a model covering only half of the target sequence is unlikely to get a score above 0.5. QMEANDisCo on the other hand evaluates the model 'as is' without explicit coverage dependency.

**Table S2. $OD_{600}$ of overnight seed cultures of test strains/reference strains expressing *ldcsm1* mutants carrying truncations of the HD loops or mutations on charged amino acids in the HD loops. Related to Fig. 1 and Fig. 3**

| $OD_{600}$ of seed / Mutation | Reference strain (CK) | NTR reference strain (CK NTR) | Test strain |
|---|---|---|---|
| WT | 3.31 | 3.1 | 3.15 |
| d-L1 | 3.17 | 3.31 | 3.2 |
| d-L2a | 3.21 | 3.23 | 3.31 |
| d-L2b | 3.09 | 3.15 | 3.1 |
| E63A | 3.15 | 3.12 | 1.84 |
| K65AK66A | 3.2 | 3.24 | 3.31 |
| R70A | 3.2 | 3.21 | 3.36 |
| E73AE76A | 3.19 | 3.14 | 3.06 |
| D94A | 3.24 | 3.2 | 3.16 |
| R95AR96A | 3.07 | 3.14 | 3.1 |
| 2E | 3.18 | 3.15 | 1.53 |
| K102A | 3.1 | 3.36 | 3.3 |
| D106A | 3.18 | 3.33 | 3.16 |
| D113A | 3.29 | 3.11 | 3.16 |

Single colony of each strain is inoculated into 5ml LB containing 100µg/ml Amp and 50µg/ml Kan. Seed cultures of each strain is cultivated at 37°C, 220 rpm for 12h.

**TableS3 ssDNA primers and substrates used in this study**

| ssDNA | Sequence (5'-3') | Description |
|---|---|---|
| **Primers** | | |
| Cas-F | TCCGCGCACATTTCCCCGAACCGCCCTGCAGATCCGGA | pColE-Cas-S1 construction |
| Cas-R | GTAGGTGTTCCACAGGGTAGAATAGGCGTATCACGAGGCCCT | pColE-Cas-S1 construction |
| ColE-F | GGCCTCGTGATACGCCTATTCTACCCTGTGGAACACCTACATC | pColE-Cas-S1 construction |
| ColE-R | TCCGGATCTGCAGGGCGGTTCGGGGAAATGTGCGCGGA | pColE-Cas-S1 construction |
| arab-F | GCTAGCAGAAACCAATTGTCCATATTGCATCAGAC | p15-G/CTR/NTR construction |
| arab-R | GAGCTCAAAAAAAAGATTTCAACAGAACCGTTTCTACTCAA TGA | p15-G/CTR/NTR construction |
| p15A-F | GATGCAATATGGACAATTGGTTTCTGCTAGCGACGAAAGGGCC T | p15-G/CTR/NTR construction |
| P15A-R | CGGTTCTGTTGAAAATCTTTTTTTTGAGCTCCGTCGACAAGCT TG | p15-G/CTR/NTR construction |
| Overlap-F | ACCCTTTATCTTTAAGCGGTAAGATTGA | LdCsm1 /Csm3 mutation |
| Overlap-F | TAAGGCCAAGCGTCAATCTTACC | LdCsm1 /Csm3 mutation |
| d-L1a-F | CTACACTGTCGCCAGAGCCGCCTTGATTATGGTGGTAATACAA GGCATCA | LdCsm1 d-L1a truncation |
| d-L1a-R | CCATAATCAAGGCGGCTCTGGCGACAGTGTAGCTTATCTGACT TATATTGGT | LdCsm1 d-L1a truncation |
| d-L1b-F | CAGATAAGCTACACTGTCCTCAGCCTGTCTCAGATTTGTTTTTT TATTC | LdCsm1 d-L1b truncation |
| d-L1b-R | AATCTGAGACAGGCTGAGGACAGTGTAGCTTATCTGACTTATA TTGG | LdCsm1 d-L1b truncation |
| d-L1c-F | CAGATAAGCTACACTGTCCTGTCTCAGATTTGTTTTTTTATTCT CCCGT | LdCsm1 d-L1c truncation |
| d-L1c-R | AAAACAAATCTGAGACAGGACAGTGTAGCTTATCTGACTTATA TTGG | LdCsm1 d-L1c truncation |
| d-L1d-F | TAAGCTACACTGTCCTGTCTTGTTTTTTTATTCTCCCGTTGATTA TGGTGG | LdCsm1 d-L1d truncation |
| d-L1d-R | AGAATAAAAAAACAAGACAGGACAGTGTAGCTTATCTGACTT ATATTGG | LdCsm1 d-L1d truncation |
| d-L2a-F | ATGGCGTTTTAGAGCCGCCAGAGCCGCCCCCAGCCGCGATGT TGTC | LdCsm1 d-L2a truncation |
| d-L2a-R | CGCGGCTGGGGGCGGCTCTGGCGGCTCTAAAACGCCATTGGC GGATATCT | LdCsm1 d-L2a truncation |
| d-L2b-F | ACCGGTTAAAGCCAGAGCCGCCCCAGCCGCCTTTGCCG | LdCsm1 d-L2b truncation |
| d-L2b-R | AGGCGGCTGGGGCGGCTCTGGCTTTAACCGGTTTGGCATCAA GCAAG | LdCsm1 d-L2b truncation |
| d-L2c-F | TTATGATCCCAGCCGCCATAGCGGCGGTCCGTCCCA | LdCsm1 d-L2c truncation |
| d-L2c-R | CGGACCGCCGCTATGGCGGCTGGGATCATAAAACGCCATTG | LdCsm1 d-L2c truncation |
| d-L2d-F | GTTTTATGATCCCAATAGCGGCGGTCCGTCCCA | LdCsm1 d-L2d truncation |
| d-L2d-R | GGACGGACCGCCGCTATTGGGATCATAAAACGCCATTG | LdCsm1 d-L2d truncation |
| E63A-F | GATTTGTTTTTTTATTCGCCCGTTGATTATGGTGGTAATACAAG | LdCsm1 E63A mutation |
| E63A-R | CCACCATAATCAACGGGCGAATAAAAAAACAAATCTGAGACA G | LdCsm1 E63A mutation |
| K65A-F | CTGTCTCAGATTTGTTTTTGCATTCTCCCGTTGATTATGGTGGT | LdCsm1 K65A mutation |

| | | |
|---|---|---|
| | AATACA | |
| K65A-R | AATCAACGGGAGAATGCAAAAACAAATCTGAGACAGGCTGA GATAC | LdCsm1 K65A mutation |
| K66A-F | CTGTCTCAGATTTGTTGCTTTATTCTCCCGTTGATTATGGTGGT AATACA | LdCsm1 K66A mutation |
| K66A-R | AATCAACGGGAGAATAAAGCAACAAATCTGAGACAGGCTGAG ATAC | LdCsm1 K66A mutation |
| K65AK66A-F | CTGTCTCAGATTTGTTGCTGCATTCTCCCGTTGATTATGGTGGT AATACA | LdCsm1 K65AK66A mutation |
| K65AK66A-R | AATCAACGGGAGAATGCAGCAACAAATCTGAGACAGGCTGAG ATAC | LdCsm1 K65AK66A mutation |
| R70A-F | TCCGGTATCTCAGCCTGTGCCAGATTTGTTTTTTATTCTCCCG TTGATTATG | LdCsm1 R70A mutation |
| R70A-R | TAAAAAAACAAATCTGGCACAGGCTGAGATACCGGAAGAC | LdCsm1 R70A mutation |
| D73AD76A-F | CTACACTGTCTGCCGGTATCGCAGCCTGTCTCAGATTTGTTTTT TTATTCTC | LdCsm1 D73AD76A mutation |
| D73AD76A-R | CTGAGACAGGCTGCGATACCGGCAGACAGTGTAGCTTATCTG ACTTATATTG | LdCsm1 D73AD76A mutation |
| D94A-F | ATTCTTCATAGCGGCGGGCCGTCCCAGCCGCGATGT | LdCsm1 D94A mutation |
| D94A-R | ATCGCGGCTGGGACGGCCCGCCGCTATGAAGAATGCG | LdCsm1 D94A mutation |
| R95AR96A-F | GCCGCATTCTTCATAGGCGGCGTCCGTCCCAGCCGCGA | LdCsm1 R95AR96A mutation |
| R95AR96A-R | GCGGCTGGGACGGACGCCGCCTATGAAGAATGCGGCAAAGGC G | LdCsm1 R95AR96A mutation |
| E98AE99A-F | GCCGCCTTTGCCGCATGCTGCATAGCGGCGGTCCGTCC | LdCsm1 E98AE99A mutation |
| E98AE99A-R | ACGGACCGCCGCTATGCAGCATGCGGCAAAGGCGGCTG | LdCsm1 E98AE99A mutation |
| K102A-F | TATGATCCCAGCCGCCTGCGCCGCATTCTTCATAGCGGC | LdCsm1 K102A mutation |
| K102A-R | CTATGAAGAATGCGGCGCAGGCGGCTGGGATCATAAAACG | LdCsm1 K102A mutation |
| D113A-F | GCCAAACCGGTTAAAGATAGCCGCCAATGGCGTTTTATGATC | LdCsm1 D113A mutation |
| D113A-R | AAACGCCATTGGCGGCTATCTTTAACCGGTTTGGCATCAAGCA | LdCsm1 D113A mutation |
| D113N-F | GCCAAACCGGTTAAAGATATTCGCCAATGGCGTTTTATGATC | LdCsm1 E640AS641K mutation |
| D113N-R | AAACGCCATTGGCGAATATCTTTAACCGGTTTGGCATCAAGCA | LdCsm1 E640AS641K mutation |
| R125A-F | CATAAGGATGCAGATAGGCTTGCCCTTGCTTGATGCCAAAC | LdCsm1 A705K mutation |
| R125A-R | CATCAAGCAAGGGCAAGCCTATCTGCATCCTTATGAAATGAAA CCGG | LdCsm1 A705K mutation |
| Csm1-seq-F | GGTAAAGCCGTCCAAAGACAGA | LdCsm1 mutation sequencing |
| Csm1-seq-R | GCTTGGCTTCCTCATATCTTGA | LdCsm1 mutation sequencing |
| **ssDNA substrates** | | |
| FAM-S10-60 | ACTATAGGGAGAATAGAATGCCCCCATTATACAATATCTACGTT TTAGATGACCCCCCCC | 5’ FAM labeled ssDNA substrate |
| FAM-16T-BHQ | TTTTTTTTTTTTTTTT | 5’ FAM labeled and 3’ |

| | | |
|---|---|---|
| 1 reporter | | BHQ1 labeled fluorescent quenched ssDNA reporter |

**Table S4. RNA substrates used in this study**

| RNA | Sequence (3’-5’) | Description |
|---|---|---|
| NTR | UGCUCUUGAAGUUUCGAAUCUAUGGGACCUCCCUUUGGUCUGAAUUGU | Target RNA with full matching 3′ anti-tag |
| PTR | AAGUUUCGAAUCUAUGGGACCUCCCUUUGGUCUGAAUUGU | Target RNA with no 3′ anti-tag |
| CTR | AAAAAAAAGUUUCGAAUCUAUGGGACCUCCCUUUGGUCUGAAUUGU | Target RNA with 6 A in 3′ anti-tag |

Full matching 3’ anti-tag of NTR is highlighted in blue, mismatching 3′anti-tag of CTR is highlighted in red.

**Table S5. Information about type III-A Csm1 used in phylogenetic analysis. Related to Fig. S7**

| | Species of type III-A Csm1 | Accession number | Length /aa | Identity to LdCsm1 /% | Pairwise distance to LdCsm1 |
|---|---|---|---|---|---|
| 1 | *Lactobacillus delbrueckii* subsp. *bulgaricus* | AQR54990.1 | 778 | 100 | 0 |
| 2 | *Lactobacillus equicursoris* | WP_009558438.1 | 781 | 59.41 | 0.431 |
| 3 | *Ligilactobacillus ruminis* | WP_195454195.1 | 760 | 51.92 | 0.534 |
| 4 | *Limosilactobacillus mucosae* | WP_242075270.1 | 758 | 46.86 | 0.64 |
| 5 | *Halolactibacillus halophilus* | WP_089831397.1 | 759 | 43.86 | 0.722 |
| 6 | *Limosilactobacillus fermentum* | WP_135252795.1 | 776 | 44.86 | 0.669 |
| 7 | *Tetragenococcus halophilus* | WP_155224396.1 | 762 | 44.33 | 0.695 |
| 8 | *Ligilactobacillus saerimneri* | WP_235720529.1 | 771 | 44.39 | 0.678 |
| 9 | *Tetragenococcus koreensis* | WP_234755460.1 | 763 | 43.89 | 0.704 |
| 10 | *Ligilactobacillus salivarius* | WP_003706181.1 | 767 | 43.49 | 0.768 |
| 11 | *Lentilactobacillus parakefiri* | WP_095354951.1 | 766 | 43.27 | 0.704 |
| 12 | *Pilibacter termitis* | WP_234984637.1 | 769 | 40.86 | 0.773 |
| 13 | *Streptococcus salivarius* | WP_247922860.1 | 756 | 39.7 | 0.787 |
| 14 | *Streptococcus suis* | NQJ22191.1 | 759 | 39.14 | 0.792 |
| 15 | *Streptococcus equinus* | ARC34389.1 | 756 | 38.81 | 0.797 |
| 16 | *Streptococcus thermophilus* | WP_014608282.1 | 758 | 38.62 | 0.782 |
| 17 | *Streptococcus sobrinus* | WP_019789096.1 | 771 | 38.13 | 0.797 |
| 18 | *Streptococcus uberis* | WP_046393285.1 | 757 | 38.6 | 0.792 |
| 19 | *Ligilactobacillus acidipiscis* | WP_056971928.1 | 737 | 38.87 | 0.937 |
| 20 | *Streptococcus mitis* | WP_033680036.1 | 746 | 36.26 | 0.852 |
| 21 | *Mycobacterium tuberculosis* | WP_057381114.1 | 812 | 34.29 | 0.894 |
| 22 | *Weissella soli* | WP_236150965.1 | 787 | 35.21 | 0.926 |
| 23 | *Mycobacterium canettii* | WP_014001446.1 | 813 | 34.45 | 0.888 |
| 24 | *Catenibacterium mitsuokai* | MCI6076816.1 | 782 | 34.9 | 0.932 |
| 25 | *Anaerobutyricum hallii* | WP_055299153.1 | 790 | 33.86 | 0.921 |
| 26 | *Anaerostipes butyraticus* | WP_201311065.1 | 784 | 33.82 | 0.926 |
| 27 | *Blautia faecicola* | WP_129257495.1 | 769 | 31.96 | 0.995 |
| 28 | *Lachnospiraceae bacterium* | MBE5871471.1 | 771 | 33.5 | 0.96 |
| 29 | *Anaerostipes hadrus* | WP_173782538.1 | 777 | 32.76 | 0.954 |
| 30 | *Lachnospira eligens* | RGZ73648.1 | 808 | 33.17 | 0.926 |
| 31 | *Dorea longicatena* | WP_055182479.1 | 767 | 32.79 | 0.989 |
| 32 | *Blautia stercoris* | WP_187558712.1 | 794 | 32.15 | 1.001 |
| 33 | *Roseburia faecis* | WP_195965881.1 | 771 | 31.38 | 0.983 |
| 34 | *Butyrivibrio crossotus* | MBD9029879.1 | 768 | 32.79 | 0.954 |
| 35 | *Eubacterium ruminantium* | SEG00723.1 | 782 | 32.56 | 0.932 |
| 36 | *Lachnospira pectinoschiza* | WP_227090192.1 | 809 | 33.21 | 0.932 |
| 37 | *Coprococcus catus* | WP_215691318.1 | 786 | 32.41 | 0.954 |
| 38 | *Butyribacter intestini* | KQC84614.1 | 796 | 32.25 | 0.949 |
| 39 | *Roseburia intestinalis* | WP_118591946.1 | 769 | 31.82 | 1.031 |
| 40 | *Roseburia hominis* | MCQ5164598.1 | 796 | 32.09 | 0.949 |
| 41 | *Faecalicatena fissicatena* | WP_173866840.1 | 793 | 31.96 | 0.949 |
| 42 | *Lachnobacterium bovis* | WP_074730519.1 | 808 | 33.21 | 0.91 |
| 43 | *Blautia wexlerae* | NSK16511.1 | 796 | 31.88 | 0.943 |

| | | | | | |
|---|---|---|---|---|---|
| 44 | *Pseudoramibacter alactolyticus* | WP_006598741.1 | 803 | 30.92 | 0.983 |
| 45 | *Blautia massiliensis* | WP_243148711.1 | 791 | 31.43 | 0.96 |
| 46 | *Eubacterium ramulus* | WP_055290818.1 | 784 | 31.41 | 0.954 |
| 47 | *Ruminococcus* sp. *AM23-1LB* | WP_243101583.1 | 794 | 31.88 | 0.943 |
| 48 | *Jingyaoa shaoxingensis* | WP_249306873.1 | 785 | 30.69 | 0.966 |
| 49 | *Staphylococcus argenteus* | WP_088811762.1 | 757 | 33.25 | 0.977 |
| 50 | *Candidatus Scybalocola faecigallinarum* | HIS47777.1 | 791 | 31.81 | 0.971 |
| 51 | *Staphylococcus hominis* | WP_242230282.1 | 757 | 33 | 0.977 |
| 52 | *Staphylococcus haemolyticus* | WP_180554307.1 | 757 | 33.12 | 0.971 |
| 53 | *Staphylococcus lugdunensis* | WP_241435022.1 | 757 | 33.79 | 0.96 |
| 54 | *Cloacibacillus evryensis* | WP_008711417.1 | 796 | 32.44 | 0.966 |
| 55 | *Staphylococcus arlettae* | WP_103388693.1 | 757 | 33.25 | 0.977 |
| 56 | *Staphylococcus pseudintermedius* | WP_100002478.1 | 757 | 33 | 0.977 |
| 57 | *Staphylococcus carnosus* | WP_053464995.1 | 757 | 33.12 | 0.977 |
| 58 | *Staphylococcus capitis* | WP_058586756.1 | 757 | 33 | 0.977 |
| 59 | *Mammaliicoccus sciuri* | WP_233695191.1 | 757 | 33.12 | 0.977 |
| 60 | *Staphylococcus aureus* | WP_111031456.1 | 757 | 33 | 0.977 |
| 61 | *Staphylococcus epidermidis* | WP_002502666.1 | 757 | 32.75 | 0.995 |
| 62 | *Blautia faecis* | WP_173770591.1 | 778 | 30.76 | 0.966 |
| 63 | *Olegusella massiliensis* | WP_082911207.1 | 783 | 33.33 | 0.937 |
| 64 | *Staphylococcus intermedius* | WP_096559504.1 | 757 | 31.34 | 1.043 |
| 65 | *Staphylococcus auricularis* | WP_213562820.1 | 757 | 33.17 | 0.983 |
| 66 | *Ruminococcus albus* | WP_051506446.1 | 783 | 32 | 0.977 |
| 67 | *Pseudoclostridium thermosuccinogenes* | WP_161950221.1 | 804 | 31.2 | 0.954 |
| 68 | *Peptoniphilus indolicus* | WP_004819772.1 | 784 | 32.14 | 1.007 |
| 69 | *Mammaliicoccus sciuri* | WP_107573069.1 | 757 | 32.92 | 1.001 |
| 70 | *Staphylococcus delphini* | WP_155260741.1 | 757 | 31.09 | 1.05 |
| 71 | *Syntrophomonas palmitatica* | WP_172674216.1 | 794 | 31.62 | 0.989 |
| 72 | *Zhenpiania hominis* | WP_187302815.1 | 778 | 30.28 | 0.989 |
| 73 | *Massilistercora timonensis* | WP_106788669.1 | 770 | 31.61 | 0.995 |
| 74 | *Enterococcus italicus* | WP_007208958.1 | 755 | 31.37 | 0.983 |
| 75 | *Lactococcus raffinolactis* | WP_138492543.1 | 757 | 32.47 | 0.977 |
| 76 | *Mogibacterium timidum* | WP_207718830.1 | 773 | 31.54 | 0.971 |
| 77 | *Mammaliicoccus lentus* | WP_194200259.1 | 757 | 33.25 | 1.001 |
| 78 | *Syntrophomonas wolfei* | WP_011641150.1 | 808 | 31.29 | 0.995 |
| 79 | *Lactococcus lactis* subsp. *lactis* | AGA14263.1 | 757 | 32.35 | 0.983 |
| 80 | *Faecalibacterium prausnitzii* | MBS4922203.1 | 794 | 30.69 | 1.05 |
| 81 | *Butyricicoccus intestinisimiae* | WP_216468813.1 | 786 | 30.61 | 1.037 |
| 82 | *Butyricicoccus porcorum* | WP_165761179.1 | 780 | 31.07 | 1.007 |
| 83 | *Faecalibacterium hattorii* | WP_181567078.1 | 794 | 30.79 | 1.025 |
| 84 | *Anaerovibrio lipolyticus* | MCF2601464.1 | 824 | 32.1 | 0.96 |
| 85 | *Lactococcus hodotermopsidis* | WP_172207039.1 | 760 | 32.83 | 0.989 |
| 86 | *Megasphaera micronuciformis* | WP_006943178.1 | 818 | 31.4 | 0.96 |
| 87 | *Thermoclostridium caenicola* | WP_188118453.1 | 805 | 29.92 | 1.068 |
| 88 | *Phascolarctobacterium succinatutens* | MBS5425956.1 | 831 | 30.89 | 0.971 |
| 89 | *Veillonella parvula* | WP_012865002.1 | 849 | 31.14 | 0.926 |
| 90 | *Anaerovibrio slackiae* | WP_154405514.1 | 854 | 30.29 | 1.05 |

| | | | | | |
|---|---|---|---|---|---|
| 91 | *Pectinatus sottacetonis* | WP_196593208.1 | 878 | 29.71 | 0.966 |
| 92 | *Macrococcus canis* | WP_164941681.1 | 755 | 30.98 | 1.056 |
| 93 | *Enterococcus cecorum* | WP_248621361.1 | 750 | 30.75 | 1.043 |
| 94 | *Veillonella tobetsuensis* | WP_105092303.1 | 849 | 31.07 | 0.921 |
| 95 | *Faecalibacterium longum* | WP_227618859.1 | 794 | 30.48 | 1.031 |
| 96 | *Acidaminococcus fermentans* | MCF0139913.1 | 837 | 30.19 | 1.007 |
| 97 | *Megasphaera cerevisiae* | WP_048515486.1 | 837 | 29.92 | 0.971 |
| 98 | *Veillonella dispar* | WP_005384901.1 | 851 | 30.96 | 0.932 |
| 99 | *Macrococcus caseolyticus* | WP_133445583.1 | 755 | 30.82 | 1.068 |
| 100 | *Veillonella atypica* | WP_156931921.1 | 851 | 30.83 | 0.915 |
| 101 | *Megamonas funiformis* | WP_193526118.1 | 828 | 30.05 | 1.019 |
| 102 | *Megamonas rupellensis* | WP_204921187.1 | 828 | 29.85 | 1.025 |
| 103 | *Selenomonas bovis* | WP_031588317.1 | 831 | 30.37 | 1.001 |
| 104 | *Megamonas hypermegale* | WP_115151325.1 | 827 | 29.73 | 1.031 |
| 105 | *Pectinatus frisingensis* | WP_231038389.1 | 868 | 29.82 | 0.977 |
| 106 | *Anaerofustis stercorihominis* | WP_138268827.1 | 805 | 29.56 | 1.037 |
| 107 | *Megasphaera hexanoica* | WP_170087869.1 | 831 | 30.63 | 1.001 |
| 108 | *Thermosyntropha lipolytica* | WP_073093420.1 | 843 | 29.38 | 1.013 |
| 109 | *Selenomonas felix* | WP_102492655.1 | 828 | 29.82 | 1.013 |
| 110 | *Acidaminococcus provencensis* | MCH4097492.1 | 845 | 30.07 | 0.949 |
| 111 | *Mitsuokella jalaludinii* | MCI7063096.1 | 816 | 30.81 | 0.989 |
| 112 | *Anaeroglobus geminatus* | WP_216436724.1 | 750 | 32.49 | 0.995 |
| 113 | *Megasphaera elsdenii* | WP_074787444.1 | 831 | 28.17 | 1.025 |
| 114 | *Veillonella caviae* | MCI5708370.1 | 872 | 29.78 | 0.96 |
| 115 | *Koleobacter methoxysyntrophicus* | WP_241754979.1 | 840 | 30.01 | 0.966 |
| 116 | *Thermosinus carboxydivorans* | WP_007288120.1 | 838 | 29.31 | 0.995 |
| 117 | *Thermincola potens* | WP_013120872.1 | 904 | 28.68 | 0.977 |
| 118 | *Dialister invisus* | WP_227137662.1 | 851 | 29.21 | 1.037 |
| 119 | *Dialister succinatiphilus* | MCI6029968.1 | 835 | 28.02 | 1.043 |
| 120 | *Fusobacterium nucleatum* | WP_005900744.1 | 841 | 29.01 | 1.05 |
| 121 | *Fusobacterium hwasookii* | WP_220280659.1 | 842 | 29.04 | 1.068 |
| 122 | *Fusobacterium periodonticum* | WP_008820661.1 | 846 | 27.32 | 1.081 |
| 123 | *Leptotrichia wadei* | WP_147003384.1 | 883 | 28.27 | 1.043 |
| 124 | *Leptotrichia hofstadii* | WP_026746289.1 | 894 | 28.13 | 1.031 |
| 125 | *Desulfolucianica intricata* | WP_066636021.1 | 834 | 26.59 | 1.265 |
| 126 | *Desulfotomaculum copahuensis* | WP_161486076.1 | 821 | 24.37 | 1.304 |
| 127 | *Thermotoga profunda* | WP_041081283.1 | 732 | 26.61 | 1.219 |
| 128 | *Thermococcus chitonophagus* | WP_068575899.1 | 751 | 26.63 | 1.242 |
| 129 | *Thermococcus piezophilus* | WP_068663804.1 | 777 | 24.29 | 1.19 |
| 130 | *Thermococcus onnurineus* | WP_012571853.1 | 777 | 24.23 | 1.219 |
| 131 | *Archaeoglobales archaeon* | RLI85854.1 | 802 | 25.31 | 1.205 |
| 132 | *Candidatus Kuenenia stuttgartiensis* | WP_164994920.1 | 832 | 23.89 | 1.312 |
| 133 | *Pyrococcus yayanosii* | WP_013906170.1 | 779 | 26.42 | 1.242 |
| 134 | *Methanotorris igneus* | WP_013799472.1 | 835 | 25.03 | 1.197 |
| 135 | *Pyrococcus horikoshii* | WP_048053047.1 | 761 | 25.19 | 1.234 |
| 136 | *Thermococcus stetteri* | MBP1912856.1 | 785 | 25.42 | 1.219 |
| 137 | *Thermococcus radiotolerans* | WP_088866078.1 | 782 | 26.13 | 1.197 |

| | | | | | |
|---|---|---|---|---|---|
| 138 | *Thermococcus indicus* | WP_139680684.1 | 779 | 25 | 1.257 |
| 139 | *Archaeoglobus fulgidus* | WP_048095157.1 | 811 | 24.43 | 1.273 |
| 140 | *Thermococci archaeon* | RLF80054.1 | 787 | 25.41 | 1.28 |
